# Gene expression evolution in pattern-triggered immunity within *Arabidopsis thaliana* and across Brassicaceae species

**DOI:** 10.1101/2020.07.29.227397

**Authors:** Thomas M. Winkelmüller, Frederickson Entila, Shajahan Anver, Anna Piasecka, Baoxing Song, Eik Dahms, Hitoshi Sakakibara, Xiangchao Gan, Karolina Kułak, Aneta Sawikowska, Paweł Krajewski, Miltos Tsiantis, Ruben Garrido-Oter, Kenji Fukushima, Paul Schulze-Lefert, Stefan Laurent, Paweł Bednarek, Kenichi Tsuda

**Author notes:** Department of Genetics, Evolution and Environment, University College London, London, United Kingdom (Shajahan Anver). Department of Computational Biology, Adam Mickiewicz University, 61-614 Poznań, Poland (Karolina Kułak). These authors equally contributed to the work (Thomas M. Winkelmüller, Frederickson Entila, Shajahan Anver). For correspondence Kenichi Tsuda.

## Abstract

Plants recognize surrounding microbes by sensing microbe-associated molecular patterns (MAMPs) to activate pattern-triggered immunity (PTI). Despite their significance for microbial control, the evolution of PTI responses remains largely uncharacterized. Employing comparative transcriptomics of six *Arabidopsis thaliana* accessions and three additional Brassicaceae species for PTI responses to the MAMP flg22, we identified a set of genes with expression changes under purifying selection in the Brassicaceae species and genes exhibiting species-specific expression signatures. Variation in flg22-triggered transcriptome and metabolome responses across Brassicaceae species was incongruent with their phylogeny while expression changes were strongly conserved within *A. thaliana*, suggesting directional selection for some species-specific gene expression. We found the enrichment of WRKY transcription factor binding sites in 5’-regulatory regions in conserved and species-specific responsive genes, linking the emergence of WRKY-binding sites with the evolution of gene responses in PTI. Our findings advance our understanding of transcriptome evolution during biotic stress.

## Introduction

The evolution of biological traits is determined both by variation in coding sequence as well as gene expression (Das Gupta and Tsiantis, 2018; Necsulea and Kaessmann, 2014). However, our understanding of heritable gene expression variation remains fragmented. The conservation of gene expression patterns over million years of evolution across species suggests a general importance of such expression patterns and indicates that they were subjected to stabilizing selection. Conversely, diversified gene expression patterns across different species may reflect neutral or adaptive evolution (Harrison et al., 2012). For example, species-specific gene expression signatures observed in human and primate neuronal tissues suggest that cognitive differences between these species might be connected to diversified expression patterns in their brains (Enard et al., 2002). However, the immanent noise in expression data complicates to distinguish between environmental and heritable effects on gene expression variation (Voelckel et al., 2017). Interpretation of gene expression evolution in samples taken from different environments can be confounded by the action of various environmental factors, such as diet, abiotic stresses and disease status, that affect gene expression (Harrison et al., 2012). Thus, the detection of heritable gene expression variation requires comparison under the same experimental conditions (Voelckel et al., 2017).

Distinguishing between different modes of evolution is crucial to understand the evolution of a given biological process. Although powerful statistical frameworks have been developed to distinguish adaptive from neutral evolution in coding sequences (Delport et al., 2009), differentiating these two modes of evolution vis-à-vis gene expression remains challenging. Attempts have been made to define a null model of neutral gene expression evolution, albeit we still lack both a consensus neutral model as well as a generally accepted statistical framework for gene expression evolution (Rohlfs et al., 2014; Khaitovich et al., 2004). An alternative way to distinguish between different modes of gene expression evolution is to employ empirical approaches to identify gene expression patterns both within and across species. For instance, if gene regulation evolves under stabilizing selection, a conserved gene expression pattern is expected both within and among species. By contrast, if gene regulation is under directional selection in a species, a distinct, highly conserved gene expression pattern within the species is expected (Romero et al., 2012; Harrison et al., 2012).

In previous comparative transcriptome studies in plants, it was noted that variation in gene expression between or within species was substantially enriched for stress-responsive genes, suggesting an important role for stress-responsive gene expression changes in adaptation to the environment (Koenig et al., 2013; Voelckel et al., 2017; Groen et al., 2020). Despite this notion, we poorly understand how stress-induced transcriptomic changes evolved in plants. However, studies comparing expression variation within and across species in unified experimental setups are rare in general and have not been described for plants.

In recent years, several close and distant relatives of the model plant *Arabidopsis thaliana*, belonging to the Brassicaceae plant family, have been genome-sequenced and used as model systems to understand the evolution of various biological traits. For instance, the comparison of the recently sequenced *Cardamine hirsuta* genome with that of *A. thaliana* has advanced our understanding of the molecular mechanisms that mediate the evolution of leaf shapes and pod shattering (Vlad et al., 2014; Gan et al., 2016; Das Gupta and Tsiantis, 2018). The genome sequences of other Brassicaceae species including *Capsella rubella* and *Eutrema salsugineum* have been used to analyse the mechanisms underlying selfing and abiotic stress-tolerance, respectively (Wu et al., 2012; Yang et al., 2013; Slotte et al., 2013). Availability of the rich genomic resources, the different degrees of phylogenetic distance to *A. thaliana*, and the feasibility to grow these Brassicaceae species in the same experimental conditions make them an excellent system for comparative genomics, transcriptomics and metabolomics.

In nature, plants are surrounded by microbes which can be potentially beneficial or pathogenic (Fitzpatrick et al., 2020). To properly respond to the presence of these microbes, plants have evolved cell-surface localized pattern recognition receptors (PRRs) that sense conserved microbe-associated molecular patterns (MAMPs) leading to activation of pattern-triggered immunity (PTI) (Albert et al., 2020; Zhou and Zhang, 2020). The two best characterized MAMPs are the bacteria-derived oligopeptides flg22 and elf18, which are sensed by their corresponding leucine-rich repeat PRRs FLAGELLIN SENSING 2 (FLS2) and EF-TU RECEPTOR (EFR), respectively, in *A. thaliana*. Treatment of plants with flg22 or elf18 elicits a set of temporally coordinated responses including rapid MAP kinase (MAPK) phosphorylation, genome-wide transcriptional reprogramming, phytohormone and secondary metabolite production, followed by inhibition of plant growth and increased resistance against pathogens (Albert et al., 2020). Although it has been mainly studied in the context of plant-pathogen interactions, PTI has recently also been implicated in the assembly of the plant microbiota, a diverse set of microbes that colonize the healthy plant (Hacquard et al., 2017; Chen et al., 2020). Thus, PTI serves as the key mechanism that allows plants to adapt to different environments characterized by different microbial communities.

Despite the significance of PTI for plant adaptation to the environment, our understanding of PTI evolution is limited to the evolution of PRRs. For instance, *FLS2* is conserved throughout many plant lineages including Brassicaceae, Solanaceae and Poaceae families, whereas *EFR* appears to be restricted to the Brassicaceae family (Boutrot and Zipfel, 2017). However, conservation of PTI responses among different species and how PTI responses evolve remain poorly understood. Here, we took a comparative transcriptomic and metabolomic approach using *A. thaliana* (six accessions), *C. rubella*, *C. hirsuta*, and *E. salsugineum* in a unified experimental setup with multiple time points to address the evolution of flg22-triggered responses in plants.

## RESULTS

### The tested Brassicaceae plants respond to the MAMP flg22

Based on our analysis using TIMETREE (see Methods), the Brassicaceae species *C. rubella*, *C. hirsuta*, and *E. salsugineum* diverged from *A. thaliana* approximately 9, 17 and 26 Mya ago, respectively (Figure 1A). We first investigated whether flg22 treatment induces rapid phosphorylation of MPK3 and MPK6, a typical early event during PTI, in these four Brassicaceae plants. Although protein extracts from *C. hirsuta*, including those from the Oxford accession, were described to not bind flg22 (implying that *C. hirsuta* does not sense flg22) (Vetter et al., 2012), we observed a clear phosphorylation of MPK3 and MPK6 upon flg22 treatment in all tested Brassicaceae plants including *C. hirsuta* Oxford, which was absent in the *A. thaliana fls2* mutant (Figure 1B). We also observed induction of *WKRY29*, a widely used immune marker gene in *A. thaliana* (Asai et al., 2002), in all tested species at 1, 9, and 24 h after flg22 application (Figure 1C). Thus, all four tested Brassicaceae species sense flg22 to trigger typical early PTI responses as observed in *A. thaliana*.

**Figure 1.**
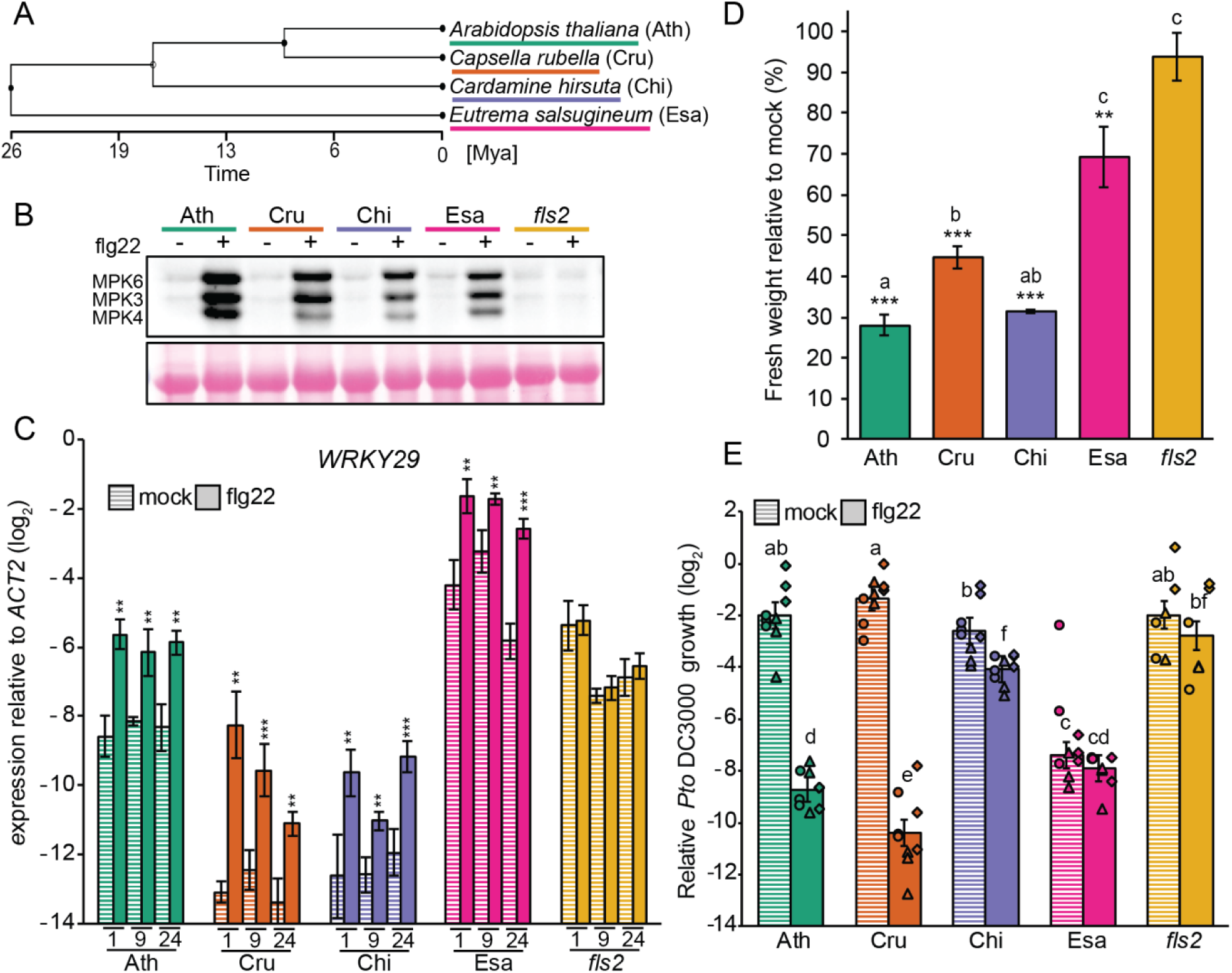
All tested Brassicaceae species sense flg22. (**A)** Phylogenetic tree generated with TimeTree (www.timetree.org) indicating the evolutionary distances between the 4 Brassicaceae species used in this study. Mya, million years ago. Ath, *A. thaliana* (Col-0); Cru, *C. rubella* (N22697); Chi, *C. hirsuta* (Oxford); Esa, *E. salsugineum* (Shandong). (**B**) 12-day-old seedlings were treated with mock or 1 µM flg22 for 15 min, and MAPK phosphorylation was measured by immunoblotting using an antiP42/44 antibody. Ponceau staining is shown as a loading control. Experiments were repeated at least three times with similar results. (**C**) Expression of *WRKY29* was determined by RT-qPCR at 1, 9, and 24 h after mock or 1 µM flg22 treatment of 12-day-old seedlings. Bars represent means and SEs of log_2_ expression levels relative to *ACTIN2* calculated from 3 independent experiments. Asterisks indicate significant difference to mock (mixed linear model followed by Student’s t-test, **, p < 0.01; ***, p < 0.001). (**D**) 7-day-old seedlings were transferred into liquid medium containing mock or 1 µM flg22 for 12 days. The fresh weight of 12 pooled seedlings was measured. The bars represent means and SEs from 3 independent experiments. Relative fresh weight (%) of flg22-treated seedlings compared to mock seedlings is shown. Statistical analysis was performed with log_2_-transformed raw fresh weight values. Asterisks indicate significant flg22 effects in each genotype (mixed linear model followed by Student’s t-test, **, p < 0.01; ***, p < 0.001). Different letters indicate significant differences in flg22 effects between different genotypes (mixed linear model, adjusted p < 0.01). (**E**) 5-week-old plants were syringe-infiltrated with 1 μM flg22 or mock 24 h prior to infiltration with *Pto* DC3000 (OD_600_ = 0.0002). The bacterial titre was determined 48 h after bacterial infiltration. The log_2_ ratio of copy numbers of a bacterial gene (*oprF*) and a plant gene (*ACTIN2*) was determined by qPCR and used as the relative *Pto* DC3000 growth. Bars represent means and SEs from 3 independent experiments each with 3 biological replicates (n = 9). The biological replicates from 3 independent experiments are represented by dots, triangles and squares. Different letters indicate statistically significant differences (mixed linear model, adjusted p < 0.01).

PTI activation reduces plant growth, a late PTI response detectable days after MAMP perception, which is another common measure of PTI outputs in *A. thaliana* (Gómez-Gómez et al., 1999). With the exception of the *fls2* mutant, chronic flg22 exposure reduced seedling growth in all tested species, but the extent of flg22-triggered growth reduction varied and was significantly weaker in *E. salsugineum* compared to the other three species (Figure 1D).

Another PTI output is enhanced pathogen resistance that is induced by MAMP pre-treatment of plants. For example, flg22 pre-treatment reduces proliferation of the foliar bacterial pathogen *Pseudomonas syringae* pv. *tomato* DC3000 (*Pto* DC3000) in *A. thaliana* leaves (Zipfel et al., 2004; Tsuda et al., 2008). We found that flg22 pre-treatment reduced bacterial proliferation in *A. thaliana* and *C. rubella* (Figure 1E). In contrast, *Pto* DC3000 growth was only slightly reduced in *C. hirsuta* and not altered in *E. salsugineum* by flg22 treatment (Figure 1E). Thus, the robust induction of early PTI responses by flg22 observed in all tested Brassicaceae does not necessarily lead to heightened immunity against this bacterial pathogen (Figure 1B, C). We noticed that the *Pto* DC3000 titre was much lower in *E. salsugineum* compared to the other species (Figure 1E) and speculated that type III effector(s) from *Pto* DC3000 may be recognized in *E. salsugineum*, triggering ETI and masking the flg22-triggered PTI effect. However, flg22 pre-treatment followed by inoculation with a *Pto* DC3000 mutant strain lacking the functional type III secretion system (*Pto hrcC*) revealed reduced bacterial growth in *A. thaliana* but not in *E. salsugineum* (Supplemental Figure 1). Thus, flg22 pre-treatment was ineffective against this bacterial pathogen in *E. salsugineum*. Interestingly, *Pto hrcC* did not grow in *C. rubella* (Supplemental Figure 1), suggesting that *Pto hrcC*-triggered PTI responses sufficiently limited *Pto hrcC* growth in *C. rubella*. In summary, while flg22 triggers typical PTI responses in all tested Brassicaceae plants, the physiological consequences, such as plant growth inhibition and bacterial resistance, vary across species.

### Flg22 triggers extensive transcriptional reprogramming in all tested Brassicaceae species

To study the transcriptome evolution of PTI responses, we generated RNA-sequencing (RNA-seq) data for early (1 h), intermediate (9 h), and late (24 h) transcriptome responses after flg22 or mock treatment of the four Brassicaceae species (Figure 2A). In total, this dataset comprised 72 samples with 33.3 million 100-bp strand-specific reads per sample on average. Normalised and log_2_-transformed count data were used for statistical analysis using a linear model (see Methods).

**Figure 2.**
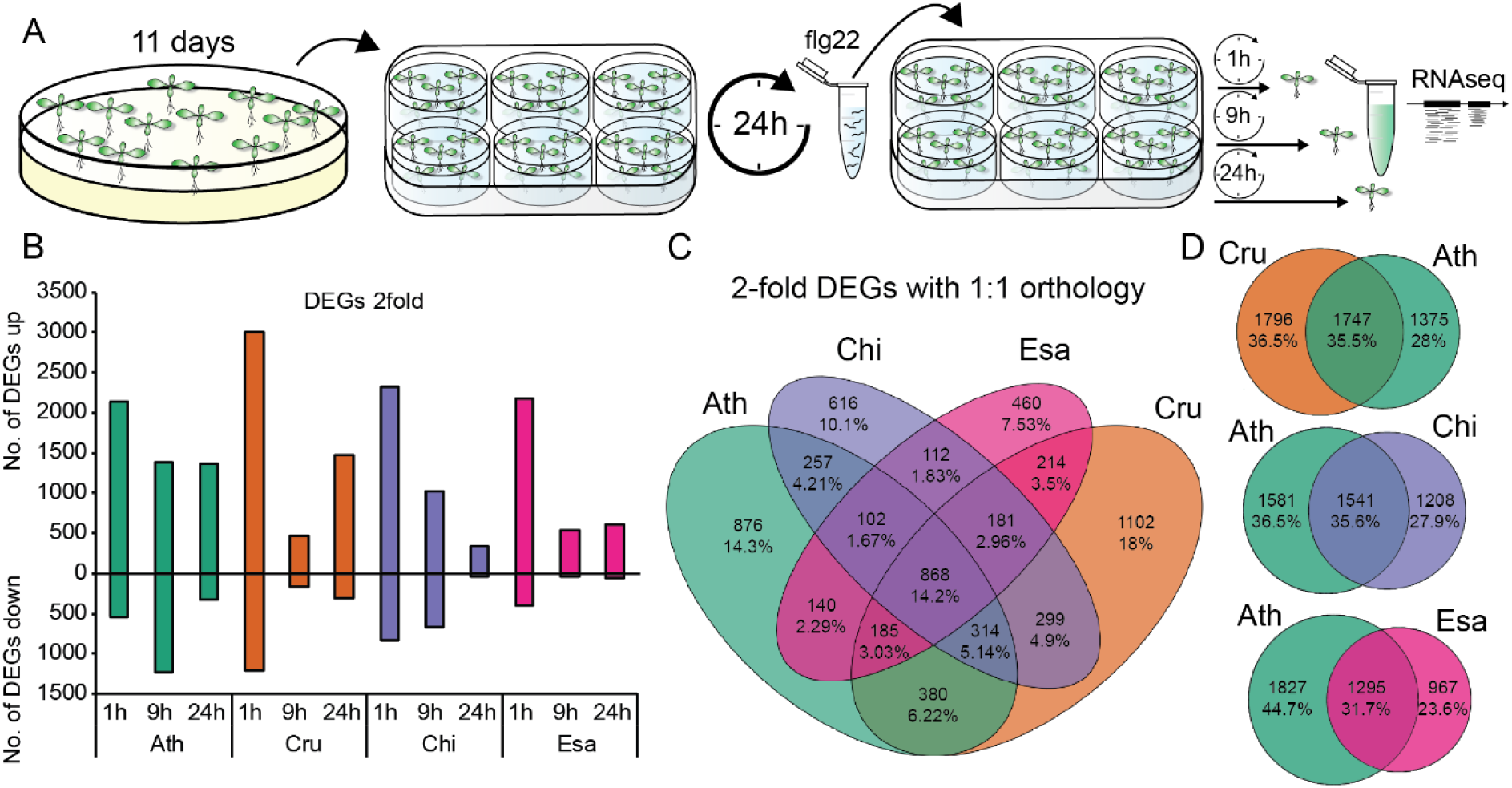
All tested Brassicaceae species trigger massive transcriptional reprogramming upon flg22 perception. (**A**) Schematic representation of the experimental design. (**B**) The number of differentially expressed genes (DEGs, q-value < 0.01 and |log_2_ fold change| > 1) for up- or down-regulated genes was plotted at the indicated time points for each species. (**C**) A Venn diagram showing shared and specific DEGs between species. All DEGs differentially expressed at least at one time point in 1 species and which showed 1:1 orthology were used. (**D**) Venn diagrams showing shared DEGs between *A. thaliana* and the indicated species. Ath, *A. thaliana* (Col-0); Cru, *C. rubella* (N22697); Chi, *C. hirsuta* (Oxford); Esa, *E. salsugineum* (Shandong).

We determined differentially expressed genes (DEGs) upon flg22 treatment compared to the mock samples based on an adjusted P-value below 0.01 and a minimum fold-change of two for each species at each time point. We observed massive transcriptional reprogramming in all species with 4,349, 4,964, 4,038, and 2,861 DEGs in *A. thaliana* (Ath), *C. rubella* (Cru), *C. hirsuta* (Chi), and *E. salsugineum* (Esa), respectively (Figure 2B). The number of upregulated genes at 1 h was comparable among species, while the number of downregulated genes at 1 h was more variable, with *C. rubella* downregulating approximately three times as many genes as *E. salsugineum*. Interestingly, the number of DEGs at later time-points differed markedly among these species: the expression of about 2,000 genes was altered in *A. thaliana* and *C. rubella*, whereas only 300 to 500 genes were differentially regulated in *C. hirsuta* and *E. salsugineum* 24 h after flg22 treatment (Figure 2B).

To allow comparison of expression changes of individual gene between species, we used Best Reciprocal BLAST to determine 1:1 orthologues between *A. thaliana* and the other Brassicaceae species. Subsequently, we only selected genes showing a 1:1 orthologue relationship between *A. thaliana* and each of the Brassicaceae species, resulting in a set of 17,856 orthologous genes (Supplemental Data Set 7). From the total of 6,106 genes, which were differentially expressed at least at one time point in at least one of the species, 868 DEGs (14.2%) were shared among all Brassicaceae species (Figure 2C and Supplemental Figure 2). These 868 DEGs represent a core set of flg22-responsive genes in these Brassicaceae species as their responses to flg22 were maintained over 26 million years of evolution. We also found that a substantial number of DEGs were species-specific (Figure 2C). The specific up- or down-regulation of 460 to 1,102 DEGs suggests substantial diversification of flg22-triggered transcriptional responses during Brassicaceae evolution. Comparisons between *A. thaliana* and each of the species revealed that about one third of flg22-induced transcriptional changes (35.5% with *C. rubella*, 35.6% with *C. hirsuta* and 31.7% with *E. salsugineum*) are shared between *A. thaliana* and each of the respective species (Figure 2D). Taken together, flg22 triggers overlapping but distinct massive transcriptional reprogramming patterns in these Brassicaceae species.

### Conserved flg22-responsive genes during Brassicaceae evolution

Next, we examined the expression dynamics of the shared 868 DEGs (Figure 3A). These shared DEGs exhibit similar expression patterns among all four species: genes induced in one species were also induced in the other species. Comparisons with publicly available datasets revealed that these shared genes are commonly responsive to MAMPs (flg22, elf18, and oligogalacturonides (OGs)) and Damage-associated molecular patterns (DAMPs; Pep2) in *A. thaliana* (Figure 3A). Many well-known genes involved in different aspects of plant immunity such as MAMP perception (*CERK1, BAK1, BIK1,* and *SOBIR1*), production of reactive oxygen species (*RBOHD*), MAPK cascades (*MKK4* and *MPK3*), salicylic acid (SA) signalling (*CBP60G, NPR1,* and *NPR3*), and immune-related transcription factors (*WRKY13/33/40/62, ERF6/104,* and *MYB51/122*) were among the conserved flg22-responsive genes (Figure 3B). In addition, a large number of genes, i.e., approximately 50% of the top 25 induced genes, whose high induction by flg22 was conserved among all tested Brassicaceae are either functionally unannotated or were not previously associated with immunity (Figure 3B; red boxes). Thus, various genes with potentially important and conserved functions in plant immunity remain to be characterized. Taken together, our data define a core set of genes with conserved flg22 responsiveness over 26 million years of Brassicaceae evolution, suggesting that the regulation of these genes is under purifying selection and therefore might be broadly relevant for plant-bacterial interactions.

**Figure 3.**
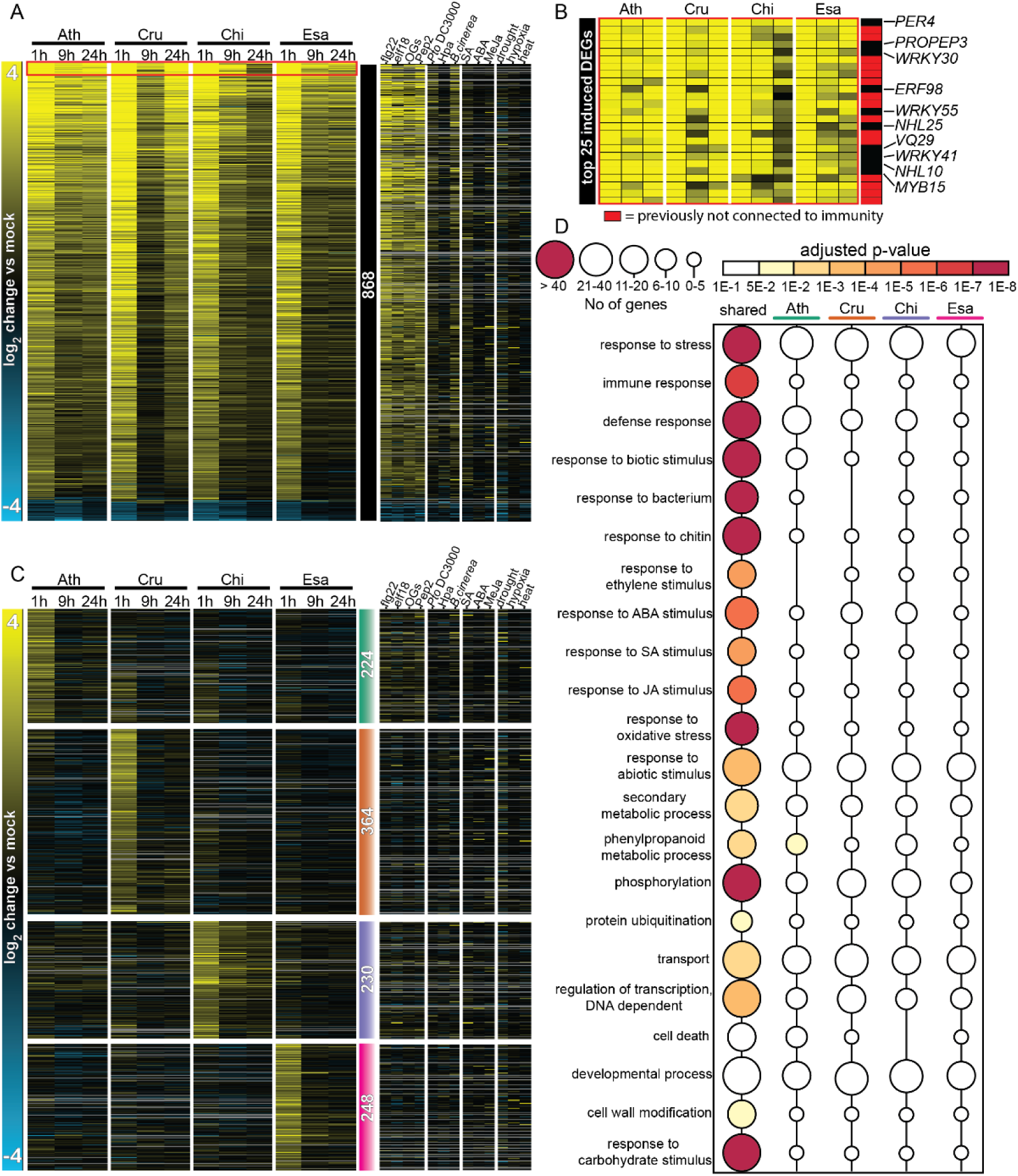
Conserved yet distinct transcriptomic responses to flg22 in Brassicaceae species. (**A**) Heatmap of 868 DEGs shared among the four tested Brassicaceae species (see Figure 2C) sorted by mean expression values. The heatmap on the right displays expression changes of the 868 DEGs under indicated stress conditions in publicly available *A. thaliana* datasets (Genevestigator). Ath, *A. thaliana* (Col-0); Cru, *C. rubella* (N22697); Chi, *C. hirsuta* (Oxford); Esa, *E. salsugineum* (Shandong). See Supplemental Data Set 1 for list of individual genes. (**B**) Heatmap of top 25 flg22-induced genes based on the mean induction of all samples. Red indicates DEGs that previously have not been implicated in plant immunity. (**C**) All 6,106 DEGs were clustered by k-means (k = 15) and 4 clusters exhibiting species-specific expression signatures are shown (see Supplemental Data Set 2). Coloured bars with the number of genes indicate Ath- (green), Cru- (orange), Chi- (purple), and Esa- (magenta) specific clusters. The right heatmap displays expression changes of these genes under indicated stress conditions in publicly available *A. thaliana* datasets (Genevestigator). See Supplemental Data Set 2 for list of individual genes. (**D**) Enrichment of selected GO terms among common DEGs and species-specific expression clusters (generated with BinGO). Circle sizes indicate the number of genes within each GO term and the colour of the circle indicates the adjusted p-values for the enrichment of respective GO terms. See Supplemental Data Set 3 for the full GO terms.

### Differences in flg22-triggered transcriptomic responses among Brassicaceae species

While in general a similar number of genes were differentially expressed after flg22 treatment in the tested Brassicaceae plants, there were substantial differences in temporal dynamics. For instance, transcriptional reprogramming was more transient in *E. salsugineum* compared to *A. thaliana*, and *C. rubella* showed a peculiar pattern characterized by a decrease in the number of DEGs at 9 h and an increase at 24 h (Supplemental Figure 3A). Rapid and sustained transcriptional responses were previously associated with effective bacterial resistance (Lu et al., 2009; Tsuda et al., 2013; Mine et al., 2018). Thus, the lack of flg22-triggered growth restriction of *Pto* DC3000 in *E. salsugineum* (Figure 1E and Supplemental Figure 1) might be explained by the transient nature of the transcriptional response in this species. To gain insights into biological processes associated with this expression pattern, we extracted genes that were induced at 1 h in both *A. thaliana* and *E. salsugineum* and were induced at 24 h in *A. thaliana* but not in *E. salsugineum* (Supplemental Figure 3B). By investigating publicly available gene expression datasets, we found that most of these genes were induced by SA in *A. thaliana* (Supplemental Figure 3C). Consistent with this analysis, flg22 treatment increased SA levels in *A. thaliana* but not in *E. salsugineum* (Supplemental Figure 3D). These results suggest that activated SA signalling is responsible for sustained transcriptional reprogramming in *A. thaliana*. However, flg22-induced transcriptome responses were comparable between wild-type *A. thaliana* Col-0 and the mutant deficient in *SID2*, encoding an SA biosynthesis gene responsible for increased SA accumulation in response to flg22 (Supplemental Figure 3E-H) (Hillmer et al., 2017). Thus, SA accumulation alone does not explain the distinct temporal dynamics of transcriptional reprogramming in these Brassicaceae plants.

A considerable number of genes were only differentially expressed in one of the Brassicaceae species (Figure 2C). To understand the degree of specificity in gene expression patterns in these Brassicaceae plants, we clustered and visualized expression changes of all 6,106 DEGs (Supplemental Figure 4). This analysis revealed that although most of the DEGs showed similar expression patterns, four gene clusters exhibited species-specific signatures (Figure 3C and Supplemental Figure 4). These four clusters contained 1,086 genes, representing about 18% of all DEGs. Although GO term analysis of the shared DEGs revealed a strong enrichment of defence-associated biological processes including “immune/defence response”, “response to bacterium”, and “response to ethylene/SA/JA stimulus”, there was almost a complete lack of GO term enrichment within the four gene clusters showing species-specific expression signatures (Figure 3D). We entertain the possibility that genes showing species-specific patterns are involved in a collection of biological processes. Indeed, species-specific clusters contained genes associated with stress- and/or immunity-associated GO terms (Figure 3D). In *A. thaliana*- and *C. hirsuta*-specific flg22-inducible genes, GO terms connected to secondary metabolism including “phenylpropanoid metabolic process”, “lignin metabolic process” and “coumarin metabolic process” were significantly enriched (Figure 3D and Supplemental Data Set 3). Distinct changes in the expression of these genes could result in differences in secondary metabolite production, which could in turn directly impact microbial growth and behaviour in plants (Piasecka et al., 2015; Stringlis et al., 2019).

### Flg22 transcriptome responses are highly conserved among genetically and geographically diverse *A. thaliana* accessions

The observed differences in gene expression patterns indicate diversification processes that might have occurred over 26 million years of Brassicaceae evolution. Alternatively, such variation in transcriptome responses can be generated within a single species. To address this question, we analysed the variation in flg22 responses among *A. thaliana* accessions. First, we tested the responsiveness of 24 *A. thaliana* accessions to flg22 using a MAPK phosphorylation assay. Flg22 treatment induced MAPK phosphorylation in all accessions except Cvi-0, which lacks a functional FLS2 receptor, therefore representing a natural negative control (Dunning et al., 2007) (Figure 4A). To avoid underestimation of diversity in flg22 responses within *A. thaliana*, we selected 12 out of the 24 accessions, which belong to distinct genetic groups (based on admixture groups from 1001genomes.org) and are geographically distributed over the USA, Europe, and Asia (Figure 4B). We found that all the 12 accessions induced *PROPEP3* expression 1 h after flg22 treatment (Figure 4C). We further generated and analysed the transcriptomes of five of these accessions 1 h after flg22 or mock treatment using RNA-seq. Importantly, these accessions were collected from geographically distant regions, were genetically diverse, and showed variable growth phenotypes (Figure 4B and D). We mapped the RNA-seq reads on the *A. thaliana* Col-0 reference genome and used the same set of 17,856 1:1 orthologous genes as in the comparison between Brassicaceae species to avoid overestimation of conservation, which may result from a larger number of shared genes in the comparison within *A. thaliana*.

**Figure 4.**
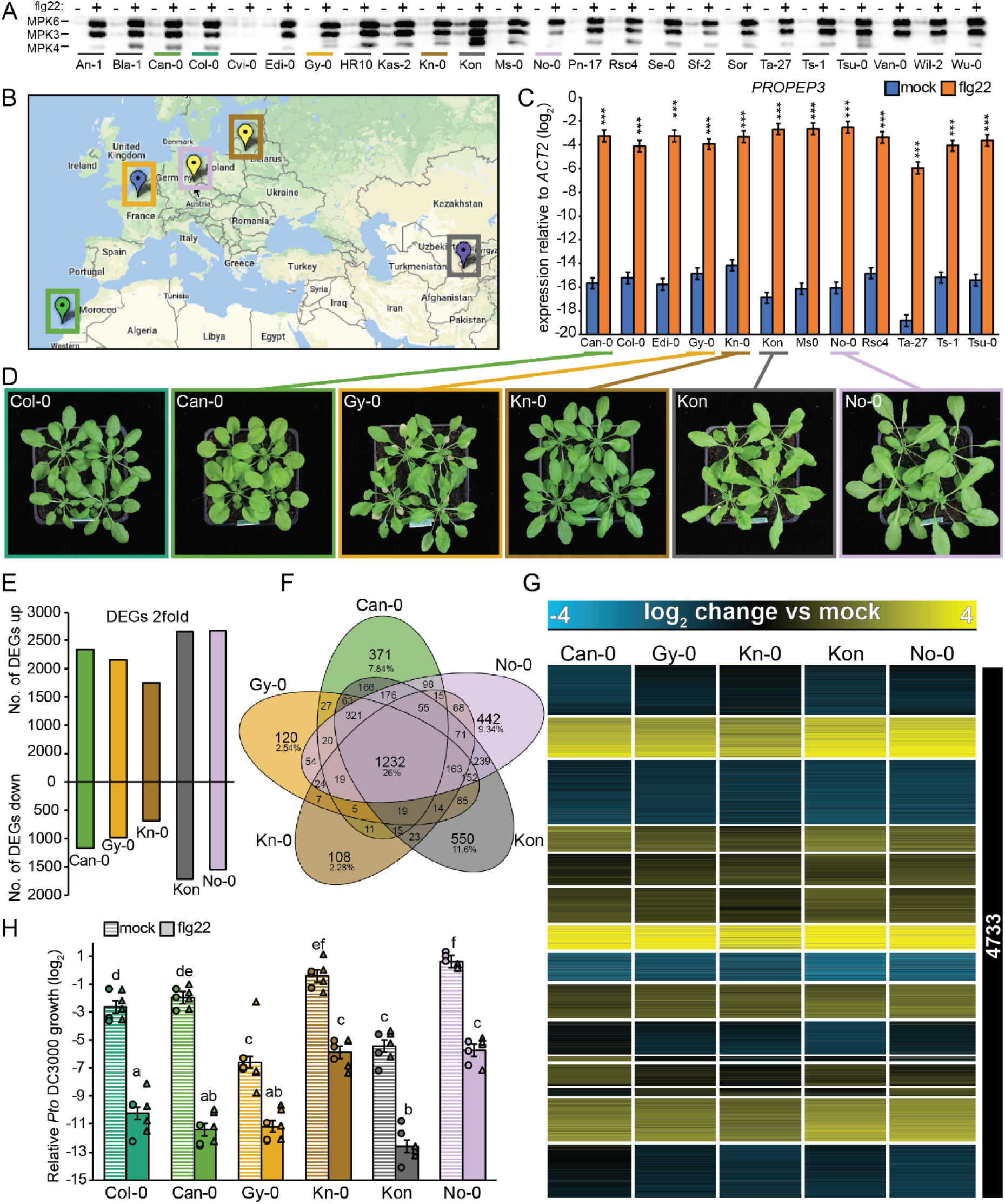
Flg22-triggered transcriptional responses show a high degree of conservation among *A. thaliana* accessions with diverse genetic backgrounds. (**A**) 12-day-old seedlings were treated with mock or 1 µM flg22 for 15 min, and MAPK phosphorylation was measured in the indicated *A. thaliana* accessions by immunoblotting using an anti-p42/44 antibody. (**B**) Geographic origins of the 5 accessions chosen for RNA-seq analysis are shown on the map created at 1001genomes.org. The colours of the markers indicate different genetic groups determined by The 1001 Genomes Consortium (1001 Genomes Consortium, 2016). (**C**) 12-day-old *A. thaliana* seedlings were treated with mock or 1 μM flg22 for 1 h, and expression of *PROPEP3* was determined by RT-qPCR. The accessions highlighted in colour were used for the RNA-seq experiments. Bars represent means and SEs of log_2_ expression levels relative to *ACTIN2* from 3 independent experiments. Asterisks indicate significant differences of flg22 compared to mock samples (Student’s t-test, ***, p < 0.001). (**D**) Representative pictures of the 4-week-old *A. thaliana* accessions chosen for RNA-seq. (**E-G**) 12-day-old *A. thaliana* seedlings were treated with mock or 1 μM flg22 for 1 h and extracted RNA was subjected to RNA-seq. The analysis was limited to the list of 17,856 genes showing 1:1 orthology in all tested Brassicaceae species to directly compare inter- and intra-species variation in transcriptome responses. DEGs were defined using the following criteria: q-value < 0.01 and |log_2_ fold change| > 1. (**E**) Bars represent the number of up- or down-regulated DEGs in each *A. thaliana* accession. (**F**) A Venn diagram showing shared and specific DEGs in *A. thaliana* accessions. (**G**) Heatmap of DEGs in at least 1 accession clustered by k-means (k = 15). Log_2_ expression changes compared to mock are shown. See Supplemental Data Set 4 for list of individual genes. (**H**) 5-week-old plants were syringe-infiltrated with mock or 1 μM flg22 24 h prior to infiltration with *Pto* DC3000 (OD_600_ = 0.0002). The log_2_ ratio of copy numbers of a bacterial gene (*oprF*) and a plant gene (*ACTIN2*) was determined by qPCR and used as the relative *Pto* DC3000 growth. Bars represent means and SEs from 2 independent experiments each with 3 biological replicates (n = 6). The biological replicates from 2 independent experiments are represented by dots and triangles. Different letters indicate significant differences (mixed linear model, adjusted p < 0.01).

The transcriptome responses of *A. thaliana* accessions to 1 h-flg22 treatment were similar in magnitude to those of other Brassicaceae plants and the *A. thaliana* Col-0 accession (4,964 to 2,861 DEGs), ranging from 4,372 (Kon) to 2,443 (Kn-0) DEGs (Figure 4E). However, the overlap of DEGs among the five *A. thaliana* accessions was greater than that of the four Brassicaceae species, as 1,232 DEGs, 26% of the total, were shared among these five accessions while 764 DEGs, 15.7% of the total at 1 h, were shared among the four Brassicaceae species (Figure 4F and Supplemental Figure 2). Consistent with this, expression patterns of all 4,733 DEGs (differentially expressed in at least one accession) were highly conserved among the five accessions, and we did not find accession-specific expression clusters with the same clustering threshold used in the interspecies comparisons (Figure 4G). Mapping the RNA-seq reads on the Col-0 reference genome potentially biased the analysis toward similar expression patterns among *A. thaliana* accessions. To test for this possibility, we generated SNP-corrected genomes for each accession and re-mapped the RNA-seq reads from *A. thaliana* accessions to their own genomes. These two mapping procedures yielded comparable results (Supplemental Figure 5). Therefore, we used the initial mapping procedure to the Col-0 reference genome for the following analyses.

Variable transcriptome responses to flg22 were associated with distinct effects of flg22 pre-treatment on *Pto* DC3000 growth in Brassicaceae plants (Figure 1E and Supplemental Figure 1). Therefore, we speculated that the high similarity in flg22-induced transcriptional reprogramming observed in *A. thaliana* accessions would lead to similar flg22 effects on flg22-triggered immunity against this bacterial pathogen. Indeed, flg22 significantly reduced *Pto* DC3000 titres in all accessions, although the bacterial growth in mock conditions differed among accessions with lower bacterial titres in Gy-0 and Kon and higher titres in No-0 compared to the Col-0 accession (Figure 4H). Taken together, these results indicate that evolution rates within *A. thaliana* were insufficient to diversify early flg22-induced gene expression changes and bacterial resistance.

### Inter-species variation in transcriptome responses to flg22 exceeds intra-species variation and is incongruent with the phylogeny

To directly compare variation in transcriptome responses to flg22 between the Brassicaceae species and within *A. thaliana*, we re-analysed the data from all 1 h samples together. We normalised, determined DEGs, and clustered log_2_ expression changes of all 5,961 DEGs. Similar to the previous analyses, the heatmap revealed gene clusters with species-specific signatures for each species but no gene cluster with *A. thaliana* accession-specific signatures (Figure 5A, B). Wrongly-assigned orthologous pairs could lead to spurious identification of species-specific expression patterns. Defining true orthologous genes between different species is challenging especially for gene families with many homologous genes. We reasoned that if the identification of genes with species-specific expression signatures resulted from the incorrect assignment of orthologues, the species-specific gene clusters should be associated with larger gene families compared to other gene clusters. However, we did not observe such a relationship (Supplemental Figure 6A). Therefore, the incorrect assignment of orthologous gene pairs unlikely explains the majority of the species-specific gene expression patterns. Another possibility is that distinct expression changes for genes showing species-specific patterns might be caused by differences in the basal expression level (in mock) among different Brassicaceae plants. Nevertheless, we did not find any consistent patterns in the basal expression levels of genes which would explain species-specific induction by flg22 (Supplemental Figure 6B). Thus, inter-species variation in transcriptome responses to flg22 among the selected Brassicaceae species clearly exceeds intra-species variation among *A. thaliana* accessions.

**Figure 5.**
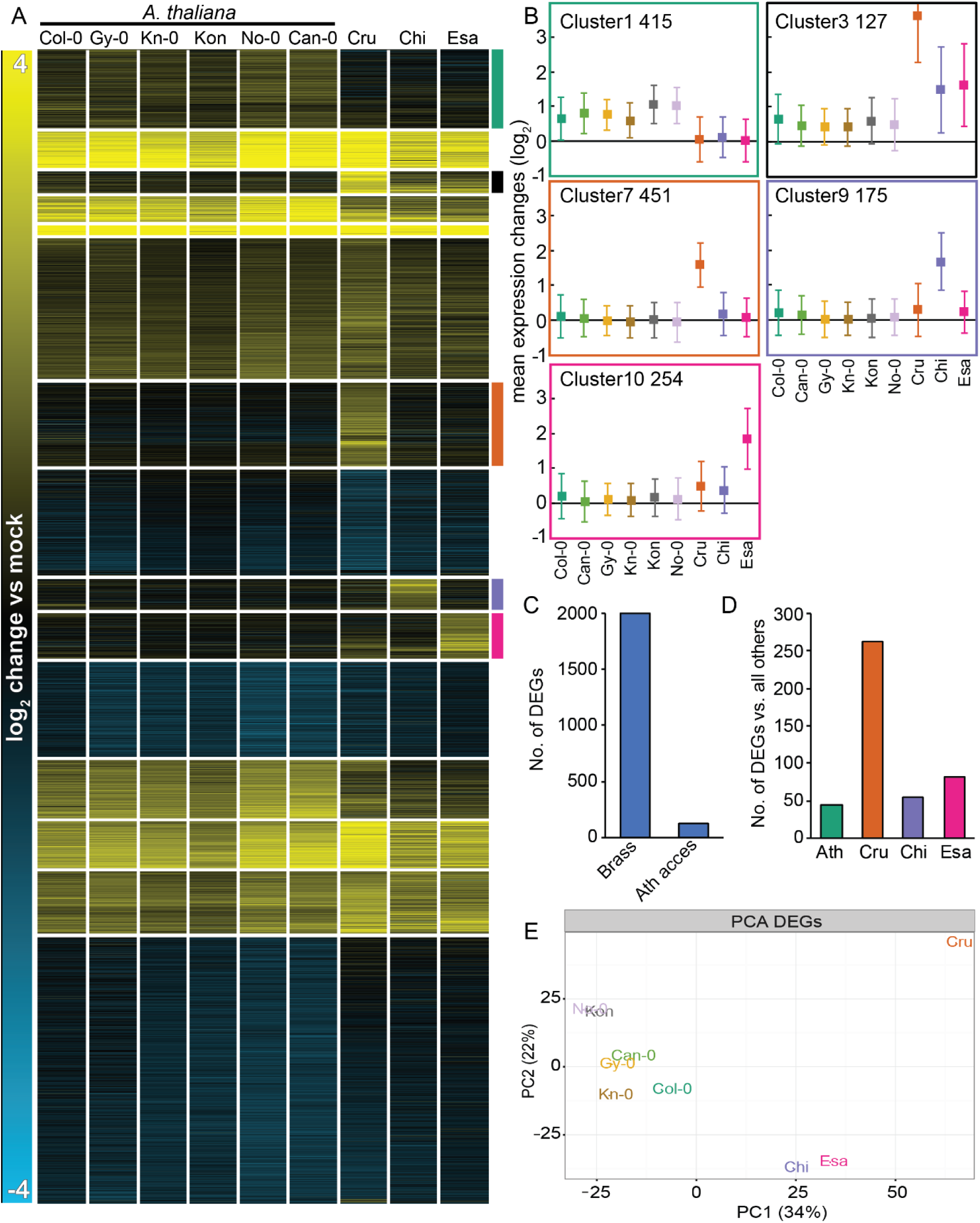
Inter-species variation exceeds intra-species variation in transcriptome responses to flg22 and is incongruent with the phylogeny. (**A**) Log_2_ expression changes of all 5,961 DEGs 1 h after 1 μM flg22 treatment were clustered using k-mean clustering (k = 15). 1:1 orthologous genes that are differentially expressed (q-value < 0.01; |log_2_ fold change| > 1) in at least 1 species or accession were used. Species-specific expression clusters are highlighted by coloured bars on the right side of the heatmap (Ath (green), non-Ath (black), Cru (orange) Chi (purple), Esa (magenta)). Cru, *C. rubella* (N22697); Chi, *C. hirsuta* (Oxford); Esa, *E. salsugineum* (Shandong). See Supplemental Data Set 5 for list of individual genes. (**B**) Mean expression changes ± SD of species-specific expression clusters in (**A**). The number of genes within each cluster is represented by the numbers on the top left side of each plot. (**C**) The total number of genes that respond to flg22 significantly differently across Brassicaceae species including *A. thaliana* Col-0 (Brass) or across *A. thaliana* accessions (Ath access). q-value < 0.01; |log_2_ fold change| > 1 criteria were used. (**D**) The number of genes that respond to flg22 significantly differently in each Brassicaceae species compared to the other 3 Brassicaceae species. q-value < 0.01; |log_2_ fold change| > 1 criteria were used. Ath, *A. thaliana* (Col-0). (**E**) Principal component analysis (PCA) of log_2_ expression changes used in (**A**).

To provide statistical support for this conclusion, we determined the number of genes that responded differently to flg22 between the Brassicaceae plants including *A. thaliana* Col-0 or between the five *A. thaliana* accessions. We detected 1,992 DEGs in the inter-species comparison and only 131 DEGs in the comparison within *A. thaliana* (Figure 5C). From these 131 genes, only the Can-0 accession harboured one gene responding differently compared to all other accessions. Among Brassicaceae plants, a considerable number of genes were specifically differentially expressed in only one of the species (Figure 5D). Consistent with *C. rubella* having the largest number of genes in the species-specific gene cluster (Figure 5A, B; Cluster 7), flg22 specifically triggered induction of the largest number of genes in *C. rubella* among the tested Brassicaceae species (Figure 5D).

The observed divergent gene expression between different species together with the low variation within species could be interpreted as evidence of species-specific directional natural selection. In other words, the species-specific gene expression signatures could be associated with adaptive evolution (Harrison et al., 2012; Romero et al., 2012). Alternatively, if the transcriptome variation among Brassicaceae species was caused by stochastic processes and was thus selectively neutral, such variation should correlate with the phylogenetic distance between the species (Broadley et al., 2008). We performed a principal component analysis (PCA) with genes that are differentially expressed 1 h after flg22 treatment in at least one plant genotype (Figure 5E). In the PCA plot, all *A. thaliana* accessions were clustered together, whereas other Brassicaceae plants were separated from *A. thaliana*. Importantly, *C. rubella*, the closest relative to *A. thaliana* among the tested species, was the most distant species from *A. thaliana* (Figure 5E). In addition, *C. hirsuta* and *E. salsugineum*, which separated 26 Mya, were clustered together (Figure 5E). Thus, the variation in flg22-induced gene expression changes across the tested Brassicaceae species was incongruent with the Brassicaceae phylogeny. These results suggest that at least some species-specific transcriptome responses to flg22 might reflect adaptive traits during Brassicaceae evolution.

The conservation of gene induction across six *A. thaliana* accessions (Figure 5A, B; Cluster 1) suggests that the observed species-specific expression signatures in Brassicaceae species might represent novel inventions in the respective species. To test this idea, we determined expression changes of selected genes showing species-specific expression signatures in different accessions or sister species of *C. rubella*, *C. hirsuta*, and *E. salsugineum* by RT-qPCR. For this, we selected *Capsella grandiflora* (Cgr, a sister species of *C. rubella*), two additional *C. hirsuta* accessions (Wa and GR2), and one additional *E. salsugineum* accession (YT). We selected *PR4*, *CYP79B2*, and *NAC32* as *C. rubella*-specific genes. *PR4* and *NAC32* were specifically induced both in *C. rubella* and *C. grandiflora*, while *CYP79B2* was induced in these two species as well as in *A. thaliana* Col-0 (Figure 6). The two *C. hirsuta*-specific genes *RAC7* and *AT3G60966* (as there is no common name for *AT3G60966*, we used the *A. thaliana* gene code) were specifically induced in all three *C. hirsuta* accessions, with the exception of *AT3G60966*, which was additionally induced in *C. grandiflora*. All three *E. salsugineum*-specific genes (*APK4*, *bZip TF*, and *CYP77A4*) were induced only in *E. salsugineum* accessions (Figure 6). Together, these findings demonstrate that the specific patterns of gene regulation observed in each of the tested Brassicaceae are conserved features in the respective species or lineage.

**Figure 6.**
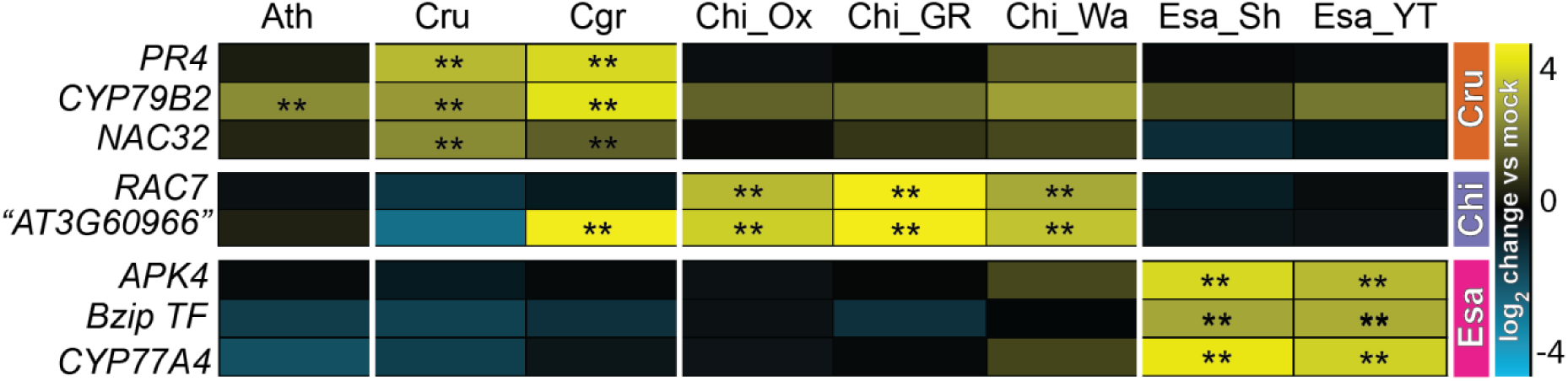
Species-specific expression signatures are conserved in sister species and accessions. Expression of selected genes showing species-specific expression signatures in Figure 5A was determined in available sister species and accessions by RT-qPCR. Gene expression was normalized to *ACTIN2*. The coloured bars on the right indicate genes showing Cru- (orange), Chi- (purple) or Esa- (magenta) specific expression signatures. The heatmap represents mean log_2_ changes of flg22 samples compared to mock from 3 independent experiments each with 2 biological replicates (n = 6). Asterisks indicate significant flg22 effects (mixed linear model, p < 0.01). Ath, *A. thaliana* Col-0; Cru, *C. rubella*; Cgr, *Capsella grandiflora*; Chi_Ox, Chi_GR, Chi_Wa, different *C. hirsuta* accessions; Esa_Sh, *E. salsugineum* Shandong; Esa_YT, Esa Yukon.

### WRKY TFs are central for flg22-triggered gene induction and may be responsible for emergence of species-specific gene induction

Changes in gene transcription are often mediated by the binding of specific transcription factors (TFs) to 5′-regulatory regions (Baxter et al., 2012). However, our understanding of how gene expression is regulated during PTI, whether gene regulatory mechanisms differ in different species, and how a given species acquires a new mode of gene regulation is far from complete. Together with genomic resources, we reasoned that our datasets, which reveal both conserved and diversified gene expression patterns in the Brassicaceae species, may provide valuable insights into these questions. To this end, we searched the 5′-regulatory regions (500 bp upstream of the transcriptional start site) of the genes in each of the 15 gene clusters (Figure 5) for known TF-binding motifs in each Brassicaceae species. Our analysis revealed that multiple motifs, which are typically bound by WRKY TFs, are highly enriched in the 5′-regulatory regions of genes in common flg22-induced clusters such as Clusters 2, 5, 13, and 14 (Figure 7A and Supplemental Figure 7). In *A. thaliana*, WRKY TFs are well known to regulate transcriptional reprogramming during plant immunity including response to flg22 (Birkenbihl et al., 2017; Tsuda and Somssich, 2015). Also, the WRKY gene family has significantly expanded in land plants, likely required for adaptation to the terrestrial environment (One Thousand Plant Transcriptomes Initiative, 2019). Our results suggest that transcriptional induction mediated by WRKY TFs is a conserved mechanism in response to flg22 across these Brassicaceae species. In addition, *A. thaliana*, *C. rubella*, and *C. hirsuta* 5’-gene regulatory regions of flg22-induced expression clusters (Clusters 13, 6, and 14, respectively) were significantly enriched for CAMTA TF-binding motifs (Figure 7 – source data 1), which play an important role in early immune transcriptional reprogramming (Jacob et al., 2018).

**Figure 7.**
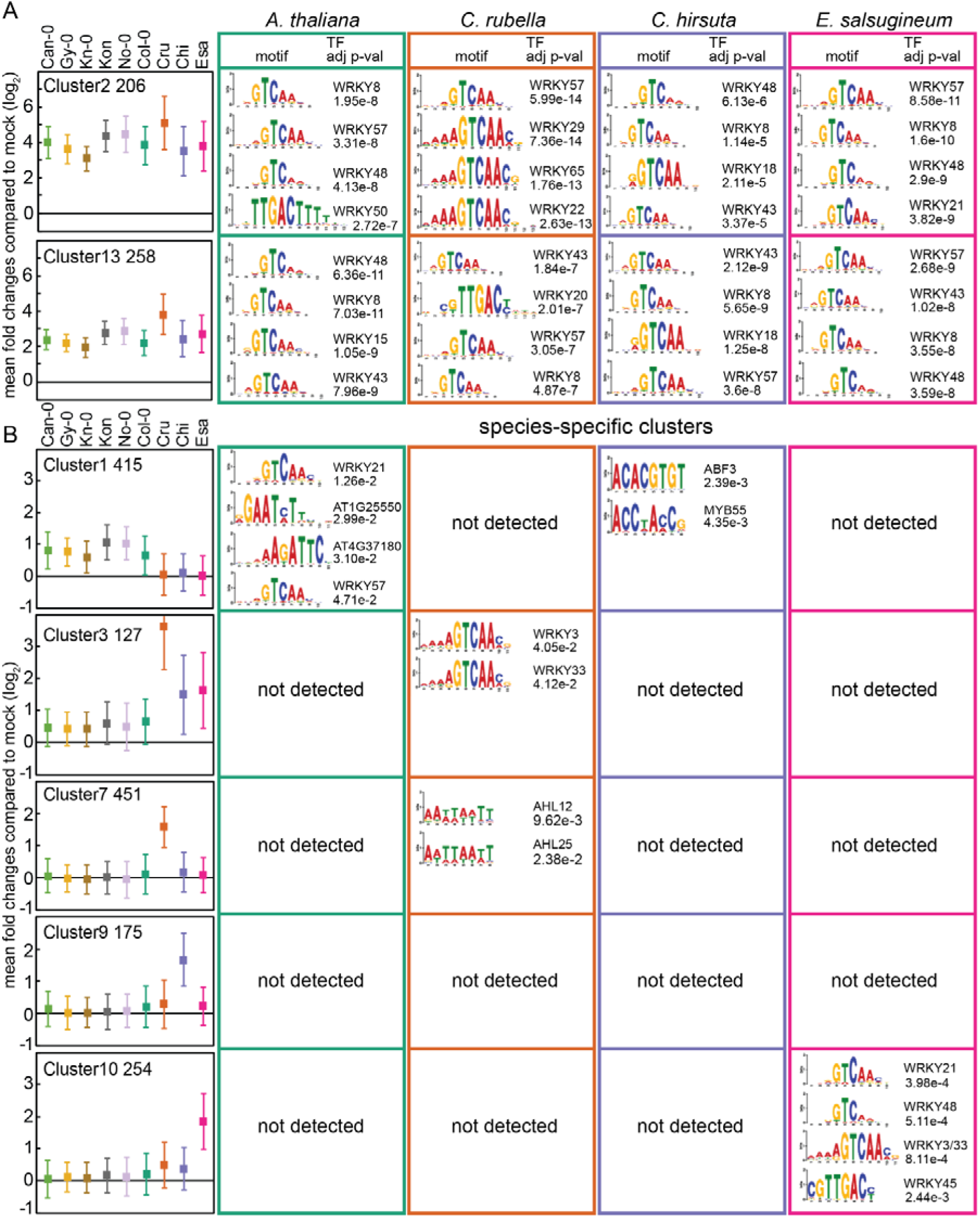
Enrichment of known TF-binding motifs in the 5′-regulatory regions of genes in shared and species-specific clusters. The 500 bp upstream sequences of the transcription start sites of the genes in the individual clusters were tested for enrichment of known TF binding motifs. Names of transcription factors, sequence logos, and adjusted p-values (up to the top 4) of motifs are shown for each Brassicaceae species. The names of clusters, the number of DEGs, and mean log_2_ fold changes ±SD compared to mock are shown on the left side. See Supplemental Data Set 6 for the other clusters. For the complete list of all enriched TF binding motifs, please see Supplemental Data Set 6. Ath, *A. thaliana* (Col-0); Cru, *C. rubella* (N22697); Chi, *C. hirsuta* (Oxford); Esa, *E. salsugineum* (Shandong).

Interestingly, we found that WRKY TFs are associated with the 5′-regulatory regions of genes showing species-specific induction only in the species that are highly flg22-responsive. For instance, in Clusters 1, 3, and 10, WRKY TF-binding motifs were only enriched in 5′-regulatory regions of flg22-induced genes in *A. thaliana*, *C. rubella*, and *E. salsugineum*, respectively (Figure 7B). In addition, in the *C. rubella*-specific expression cluster (Cluster 7), AHL TF-motifs were enriched only in *C. rubella* 5′-regulatory regions (Figure 7B). AHL TFs have been associated with plant developmental processes but some AHL TFs are involved in MAMP-induced gene expression (Lou et al., 2014; Mine et al., 2018). These results suggest that in these Brassicaceae plants the emergence of cis-regulatory sequences that are bound by specific TFs such as WRKY TFs facilitated the evolution of distinct gene induction patterns.

### Variation in coding sequences show no strong correlation with transcriptome variation

Previous studies reported a positive correlation between gene expression and coding sequence evolution and suggested that similar selective forces might have acted on both modes of evolution (Hunt et al., 2013; Khaitovich et al., 2005), although it should be noted that in some studies, this correlation was organ-dependent or not detected at all (Whittle et al., 2014; Tirosh and Barkai, 2008). Thus, the relationship between gene expression and coding sequence evolution appears to be species- or condition-dependent. Therefore, we asked whether the variation in basal or flg22-triggered expression changes is correlated with variation in amino acid sequences among the tested Brassicaceae species. We compared the standard deviation divided by the mean of the expression levels in mock-treated RNA-seq samples (1 h) of *A. thaliana* Col-0 and other Brassicaceae plants with the mean amino acid sequence identities between *A. thaliana* Col-0 and each of the other species. We found no correlation in variation between amino acid sequence and basal gene expression (Figure 8A). Similarly, we compared flg22-induced expression changes of all expressed genes or DEGs (1 h) with amino acid sequence identities and found no correlation (Figure 8B, C). Finally, we tested whether pairwise differences in flg22-induced expression changes between *A. thaliana* and individual Brassicaceae species were linked to amino acid sequence diversification. Again, we did not find any strong correlation (Figure 8D-I). Thus, variation in gene expression both at basal levels and in response to flg22 does not correlate with variation in amino acid sequences, suggesting that different selective forces drove gene expression and coding sequence evolution in these Brassicaceae species.

**Figure 8.**
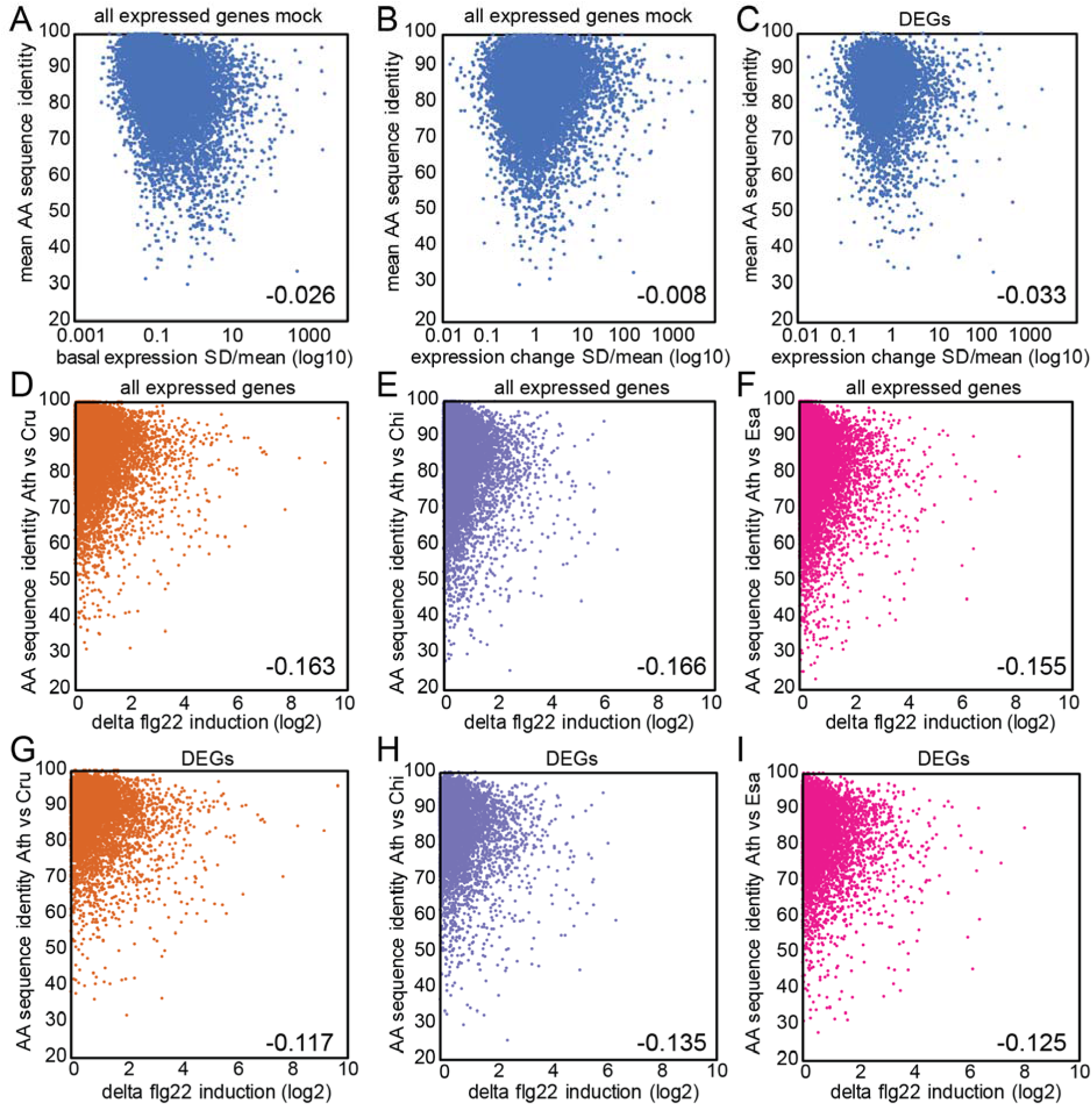
Gene expression variation does not correlate with coding sequence variation. (**A**) Mean amino acid (AA) sequence identities of *C. rubella*, *C. hirsuta* and *E. salsugineum* to *A. thaliana* (y axis) were plotted against the SD/mean of the expression values in mock samples of all four Brassicaceae plants for all expressed genes (x axis). (**B, C**) Mean AA identities of *C. rubella*, *C. hirsuta*, and *E. salsugineum* to *A. thaliana* were plotted against the SD/mean of flg22-induced expression changes in all four Brassicaceae plants for all expressed genes with 1:1 orthologs (16,100 genes) (**B**) or 5,961 DEGs (**C**). (**D–I**) Pairwise AA sequence identities of *C. rubella* (**D, G**), *C. hirsuta* (**E, H**) and *E. salsugineum* (**F, I**) to *A. thaliana* were plotted against the flg22-induced expression changes between the compared species for all expressed genes (**D–F**) or DEGs (**G–I**).

### The genetic divergence and allele frequencies of coding or regulatory regions of immediate-flg22-responsive genes implicate neutral ascendancy in *A. thaliana*

To test whether genes displaying a specific response in *A. thaliana* (clusters 1 and 12) may have been subjected to recent adaptive pressures, we compared their patterns of polymorphism and divergence at upstream and coding regions with other gene clusters displaying no *A. thaliana*-specific expression (clusters 2,5,7,9). If recent and recurrent regulatory adaptive mutations in *A. thaliana* were the ultimate cause of the expression specificity observed in clusters 1 and 12, we should observe an elevated divergence between *A. thaliana* and *A. lyrata* at regulatory regions compared to the other clusters and potentially a distribution of allele frequencies skewed towards higher frequency classes compared to neutral expectations (Nielsen, 2005). Our results, rather indicate that the genetic variation observed at genes with *A. thaliana* -specific responses are overall in line with the variation observed at other clusters regardless of species-specificity (Supplemental Figure 8). However, cluster 5 (highly induced in all species) showed the lowest genetic divergence on its upstream regions while the neutral synonymous variation for the same cluster was the highest (Supplemental Figure 8). This suggests that this lower divergence at upstream regions is not the result of a lower mutation rate but rather the result of stronger purifying selection acting on the regulatory regions of those genes with conserved expression patterns in this cluster.

The allelic distributions at the upstream and coding sequences of these clusters in *A. thaliana* were also determined to assess negative and positive selection. We did not observe clear differences in evolutionary selection among six clusters examined (Supplemental Figure 9). Long-term balancing selection has been identified as an important process shaping the genetic variation of immunity related genes in plant (Koenig et al., 2019). One of the signatures of balancing selection in polymorphism and divergence data is the increase of the ratio of non-synonymous to synonymous divergence and polymorphism (Ka/Ks and pi_a/pi_s, respectively). A number of genes under balancing selection was observed in each cluster (Supplemental Figure 10), indicating that balancing selection acts on flg22-responsive genes in *A. thaliana* independent of the species-specificity.

### Differences in metabolome profiles in response to flg22 among the Brassicaceae species

Some genes showing species-specific expression patterns were associated with GO terms connected to secondary metabolism (Supplemental Data Set 8). This prompted us to investigate whether flg22 treatment differentially affects metabolite profiles of Brassicaceae plants. In unbiased HPLC-MS analysis, we detected various differentially accumulating metabolites (DAMs; q-value < 0.05, minimum fold change of 1.5) in response to flg22 among the four Brassicaceae plants (Figure 9A). Interestingly, most flg22-induced changes in metabolite accumulation were species-specific and only 19 out of 360 DAM signals were commonly affected by flg22 in all tested Brassicaceae species, indicating a strong diversification of the native metabolome and its reprogramming in response to flg22 (Figure 9B). This notion was further supported by the clustering of log_2_ fold changes for all DAMs, which showed only limited number of overlaps between metabolome alterations (Figure 9D). In line with the flg22-triggered transcriptional activation, the variation in metabolome changes was incongruent with the Brassicaceae phylogeny as *A. thaliana* and *E. salsugineum*, which are the most distantly related tested species, clustered together in the PCA based on all DAMs (Figure 9C).

**Figure 9.**
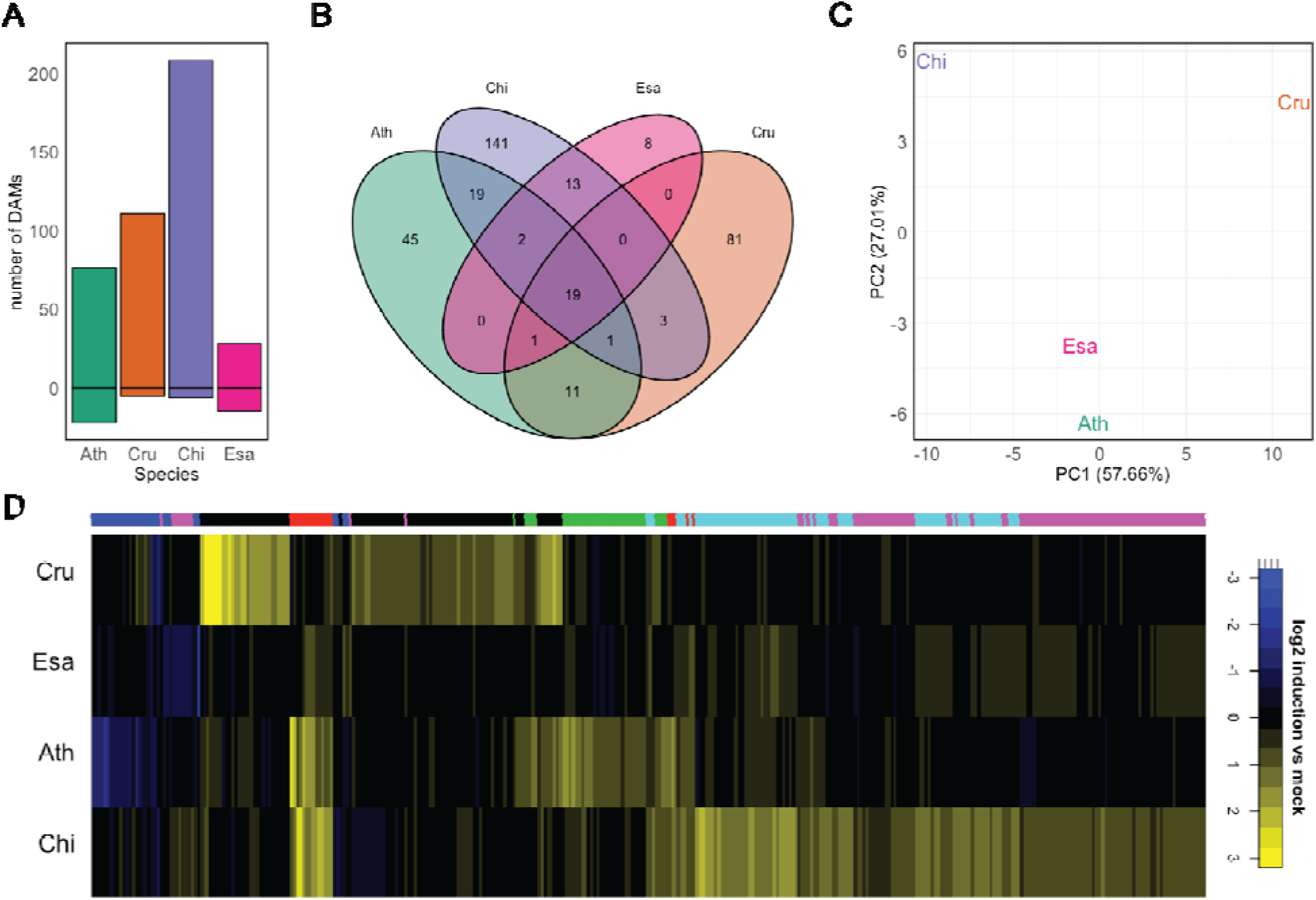
Flg22 triggers unique metabolome changes in tested Brassicaceae. Metabolite profiles were analysed with HPLC-MS 24 h after mock or flg22 treatment in 12-day-old Brassicaceae seedlings. (**A**) Differentially accumulated metabolites (DAMs) were determined using the following criteria: treatment effect or interaction treatment x species was significant with q-value < 0.05, |log_2_ fold change| > 0.585 and significance of the difference between treatment and control with p<0.05. The bars represent the number of up- or down-regulated DAMs in each species. Ath, *A. thaliana* (Col-0); Cru, *C. rubella* (N22697); Chi, *C. hirsuta* (Oxford); Esa, *E. salsugineum* (Shandong). (**B**) A Venn diagram showing shared and unique DAMs between species. All DAMs present in at least 1 species were used. (**C**) Principal component analysis of DAMs in at least 1 species. (**D**) Heatmap of log_2_ fold changes for DAMs in at least 1 species clustered by k-means clustering (k = 6; clusters marked in the color bar on top of the heatmap).

## DISCUSSION

Although current knowledge suggests that stabilizing selection and neutral fluctuation largely shape the evolution of transcriptomic variation, directional selection also appears to considerably shape the regulation of gene expression (Rifkin et al., 2003; Nuzhdin et al., 2004; Lemos et al., 2005; Whitehead and Crawford, 2006; Romero et al., 2012; Chen et al., 2019; Brawand et al., 2011). Our comparison of transcriptome responses within *A. thaliana* accessions and among four Brassicaceae species reveals fundamental features of transcriptome evolution: we have identified conserved core genes and species-specific responsive genes across Brassicaceae plants during flg22-induced PTI. A core set of responsive genes conserved across Brassicaceae likely reflect stabilizing selection to keep important gene regulatory processes in PTI whereas each species evolved their specific responses for species-specific arms. Our unified experimental setup for all transcriptome comparisons has allowed us to detect heritable gene expression variation among these species.

We have shown that transcriptome responses to flg22 are remarkably conserved within *A. thaliana*. A previous study found that variation in the expression of stress-responsive genes accounted for the majority of divergence in expression among *A. thaliana* accessions (Kawakatsu et al., 2016), and many studies have described the strong plasticity of immune-responsive gene expression in the face of other environmental perturbations such as abiotic stresses (Berens et al., 2019; Singh et al., 2014; Ueno et al., 2015; Coolen et al., 2016). Since the tested *A. thaliana* accessions were collected from habitats with distinct climatic conditions such as the Canary Islands and Lithuania and have diverse genetic backgrounds, these highly conserved early transcriptome responses to flg22 were surprising. This finding indicates that short-term evolution in divergent environments is insufficient to introduce major variation in early transcriptional responses during PTI and reflects the importance of a rapid transcriptome response in *A. thaliana* across diverse environments.

We have found that the expression changes of large numbers of genes are conserved among different Brassicaceae species during flg22-induced PTI. Many of these genes have not been previously characterized or linked to plant defence. Thus, our dataset provides the basis and rationale for future studies. In addition, the conservation of transcriptome responses to flg22 over 26 million years of Brassicaceae evolution indicates that these responses evolved under stabilizing selection (Whitehead and Crawford, 2006). Many studies have demonstrated the importance of transcription factors that regulate the expression of specific genes during PTI (Birkenbihl et al., 2017; Jacob et al., 2018). However, the lack of a method to efficiently and specifically block transcriptional reprogramming during PTI means that it remains obscure whether genome-wide transcriptional reprogramming during PTI is required for plant defence against pathogens and/or for adaptation to their environments. The stabilizing selection observed in the transcriptome responses strongly suggests that the massive transcriptional reprogramming during PTI is advantageous for Brassicaceae plants in nature.

While all tested Brassicaceae species deployed rapid, massive transcriptional reprogramming as well as MAPK activation, they exhibited different PTI outputs. For instance, in *C. hirsuta* and *E. salsugineum*, the flg22-elicited transcriptional response was not associated with flg22-induced resistance against *Pto* DC3000. This seems counterintuitive for species not benefitting from costly transcriptional reprogramming. Maintaining the PTI transcriptional reprogramming is analogous with the retention of susceptible alleles for *Rps2*, a resistance gene for the pathogen virulence effector AvrRpt2 (MacQueen et al., 2016). One of the explanations is that susceptible *Rps2* alleles encode recognition specificities for pathogen effectors that have yet to be identified (MacQueen et al., 2016). Similarly, transcriptional reprogramming induced by flg22 may be associated with effective resistance against different bacterial pathogens or the control of plant microbiota in *C. hirsuta* and *E. salsugineum* (Hacquard et al., 2017; Chen et al., 2020). Understanding of how diversity in PTI responses across plant species is linked to plant adaptation would be key to comprehend the role of PTI.

In addition to the conserved transcriptome responses, we have also revealed that each of the tested Brassicaceae species exhibits heritable species/lineage-specific expression signatures during flg22-induced PTI. This can be explained by genetic drift in which case changes might be selectively neutral. Alternatively, the species-specific expression patterns might be selective. Evolutionary theory suggests that if expression changes in a species are adaptive, little variation is expected within that species. We have shown that species-specific expression patterns are conserved among multiple accessions or sister species in the respective species. Moreover, variation in transcriptome responses during flg22-induced PTI was incongruent with the Brassicaceae phylogeny, which is inconsistent with the variation being caused solely by genetic drift. Thus, some of the species-specific expression signatures observed in this study during flg22-induced PTI appear to be selectively driven expression shifts. As similar incongruence was observed for the flg22-induced changes in metabolomes of the investigated species these expression signatures likely include genes linked with specialized metabolism.

We have also shown that interspecific differences are larger than intraspecific variations in the early transcriptome response. This expression divergence could be explained by variation in expression and function of transcription factors and/or in 5′-gene regulatory regions *cis*-regulatory elements that coincides in a lineage-specific manner. It has been thought that most of the intraspecific variations could be attributed to *cis*-associated differences which are tightly in constraint by linkage disequilibrium while interspecific differences are largely due to trans-associated alterations being a larger mutational target (Signor and Nuzhdin, 2018). However, disentangling the contributions of *trans-* and *cis-* components of transcriptional control remains challenging. A recent finding revealed that transcriptome variation is disposed to strong selection pressure under perturbed environments, in particular, genes with low expression and stochasticity, and with high plasticity (Groen et al., 2020). In line with the important role of WRKY TFs in gene induction during immunity (Tsuda and Somssich, 2015; Birkenbihl et al., 2017), we have revealed that WRKY-TF binding motifs are highly enriched in the 5′-gene regulatory sequences of species in which the genes are induced. This suggests that some specific gains of TF-binding motifs in the 5′-gene regulatory regions account for the evolution of some species-specific flg22-responsive expression changes. It might also be possible that duplicated genes could be responsible for species-specific differences that we may have missed in this study as we entirely omitted lineage-specific duplicates. However, this would not introduce a very serious bias as a large proportion of genes in the genomes (17,856) were successfully analysed in this study.

Whether gene expression evolution correlates with coding sequence evolution remains under debate (Tirosh and Barkai, 2008). Some studies found a positive correlation between gene expression and coding sequence evolution and argued that similar selection pressures act on both modes of evolution (Hunt et al., 2013; Whittle et al., 2014). In contrast, others have concluded that gene expression evolution may provide additional evolutionary capacity if the sequence of the respective gene is under evolutionary constraint (Shapiro et al., 2004; Harrison et al., 2012; Dean et al., 2015). In this scenario, gene expression variation would not be correlated with coding sequence evolution (Tirosh and Barkai, 2008; Renaut et al., 2012; Uebbing et al., 2016). In this study, we found almost no correlation between variation in basal gene expression or flg22-induced gene expression changes and variation in their amino acid sequences. The connection between gene expression and coding sequence variation might depend on species and conditions (Whittle et al., 2014). Future studies, especially in the plant field, are needed to better define the relationship between these two modes of evolution.

In summary, we have shown that variation in biotic stress-induced gene expression is under purifying selection and is also likely to be under directional selection among Brassicaceae species. Overall, our findings provide considerable insight into the evolution of transcriptome responses during stress responses in plants and constitute a basis for future research.

## Material and Methods

### Plant materials

**Table 1:** Brassicaceae species and accessions used in this study. Bold entries indicate species used for RNAseq and metabolome analyses.

| Species | Accession | Abbreviation | Reference |
| --- | --- | --- | --- |
| ***Arabidopsis thaliana*** | **Col-0** | Ath | Kenichi Tsuda lab |
| ***Capsella rubella*** | **N22697** | Cru | (Slotte et al., 2013) |
| *Capsella grandiflora* | unknown | Cgr | (Slotte et al., 2013) |
| ***Cardamine hirsuta*** | **Oxford** | Chi | (Hay and Tsiantis, 2006) |
| *Cardamine hirsuta* | Wa | Wa | (Cartolano et al., 2015) |
| *Cardamine hirsuta* | GR2 | GR | Miltos Tsiantis lab |
| ***Eutrema salsugineum*** | **Shandong** | Esa | Miltos Tsiantis lab |
| *Eutrema salsugineum* | Yukon | Eyt | Miltos Tsiantis lab |

**Table 2:**
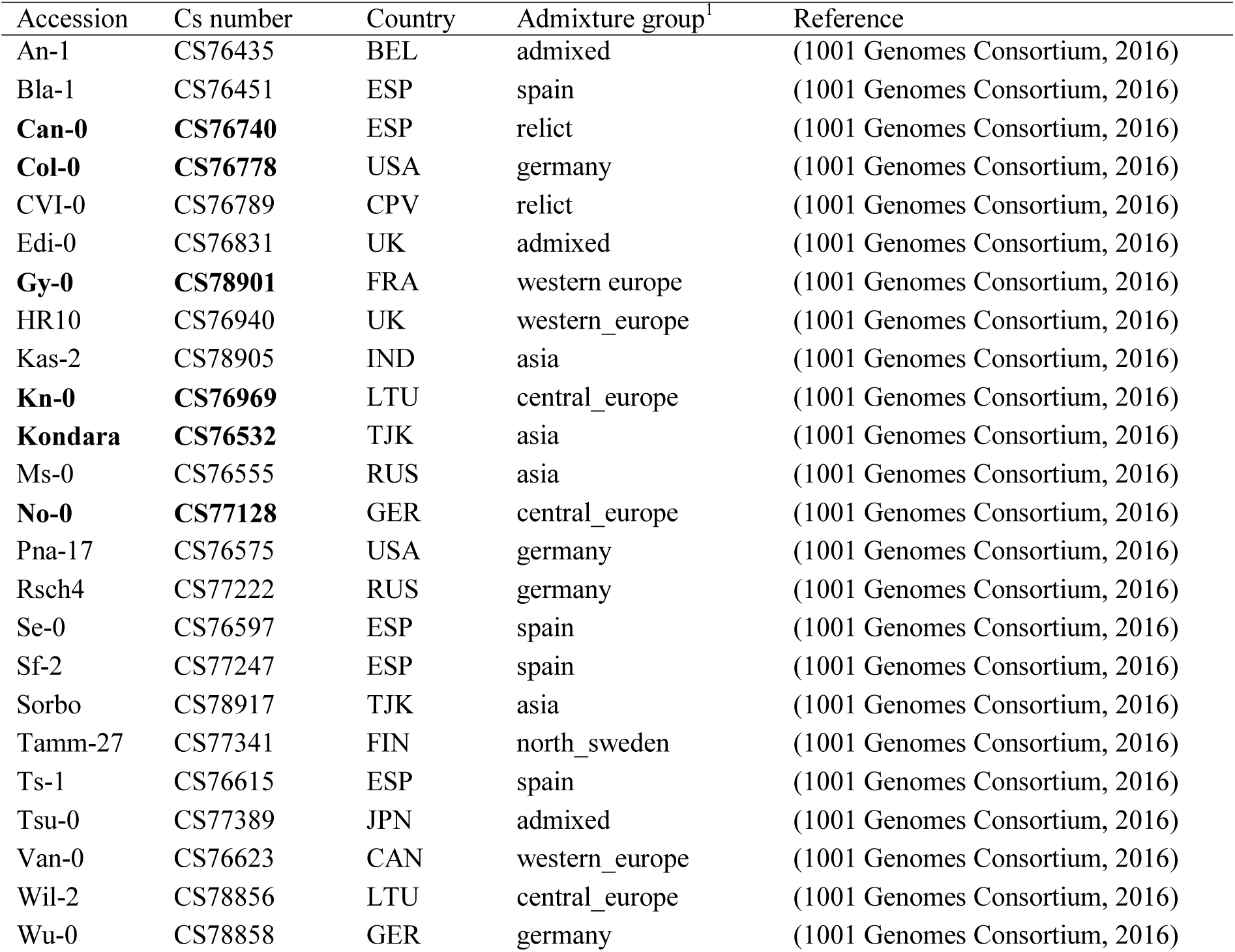
*A. thaliana* accessions used in this study. Bold entries indicate accessions used for RNAseq. ^1^Admixture group (1001 Genomes Consortium, 2016)

**Table 3:** *A. thaliana* mutants used in this study.

| Species | Mutant allele | Locus | Source |
| --- | --- | --- | --- |
| *Arabidopsis thaliana* | *sid2-2* | AT1G74710 | Tsuda et al., 2008 |
| *Arabidopsis thaliana* | *fls2* (SAIL_691C4) | AT5G46330 | Zipfel et al., 2004 |

### Plant growth

Plant seeds were sterilized by vortexing in 70% ethanol for 5 min and then 6% NaClO for 10 min, washed five times with sterile water, and stratified in sterile water at 4°C for five to seven days. Sterilized seeds were grown on ½ Murashige and Skoog (MS)-Agar (2.45 g/L M&S+Vitamins+MES (Duchefa, Netherlands), 1% sucrose, 0.5% plant agar, pH 5.8) plates in a Percival plant growth chamber (CU-36LX5D, Percival, USA) at 22°C with 10 h of light for eleven days if not stated otherwise. Eleven-day-old seedlings were transferred to liquid ½ MS-Medium (2.45 g/L M&S+Vitamins+MES (Duchefa, Netherlands), 1% sucrose) one day before flg22 treatment. Alternatively, 12-day-old seedlings were transferred to soil (Stender, Schermbeck Germany) and grown at 23°C/20 °C with 10 h/14 h (light/dark) and 60% relative humidity. Soil-grown plants were transferred to another chamber at 22°C with a 12 h photoperiod and 60% relative humidity three days before bacterial inoculation.

### Flg22 treatment

Eleven-day-old seedlings were transferred from ½ MS-Agar to 24-well plates each with 1.6 ml of ½ MS-Medium 24 h prior to treatments. If not otherwise stated, five to ten seedlings per sample were transferred to each well. For the flg22 treatment, 800 µl of 3 µM flg22 (EZBiolab Inc., USA) solution was added to the medium containing the seedlings resulting in a final concentration of 1 µM flg22. Seedlings were harvested in liquid nitrogen at the indicated time points and three wells were combined into one biological sample. The samples were stored at -80°C until use.

### Seedling growth inhibition assay

Seven-day-old seedlings grown on ½ MS-Agar were transferred to 1.6 ml of ½-MS-Medium with or without 1 µM flg22 and grown for another 12 days in these conditions. Then, the fresh weight of 12 pooled seedlings was measured. The experiment was carried out three independent times, and statistical analysis was performed with log_2_-transformed fresh weight values.

### Bacterial growth assay

For preparation of bacterial inoculum, *Pseudomonas syringae* pv. *tomato* DC3000 (*Pto* DC3000) or the T3SS deficient *Pto* DC3000 mutant *Pto hrcC* (Tsuda et al., 2008) was grown on NYGA agar (2% glycerol, 0.5% Bacto Peptone, 0.3% yeast extract, 1% Bacto Agar, pH 7.0) plates containing 25 µg/ml rifampicin for three days at 28°C. Then, bacterial strains were transferred to liquid NYGA medium containing 25 µg/ml rifampicin and incubated overnight at 28°C with shaking at 200 rpm to a final OD_600_ between 0.8 and 1. The bacteria were pelleted by centrifugation at 5,000 rpm and washed twice with sterile 5 mM MgSO_4_ before dilution to an OD600 of 0.0002 (*Pto* DC3000) or 0.001 (*Pto hrcC*).

Four to five-week-old plants were used. Two leaves per plant were infiltrated with 1 µM flg22 or sterile water (mock) using a needleless syringe. One day later, leaves treated with flg22 or mock solution were infiltrated in the early afternoon with the bacterial suspension. Two days after bacterial infiltration, two leaf disks (0.565 cm^2^) per sample from two leaves were crushed in 400 µl sterile MgSO_4_ using a Retsch mixer mill. Dilution series were made and streaked on NYGY agar plates containing 25 µg/ml rifampicin. The plates were incubated for two days at 28°C before colony forming units (cfu) were counted.

Alternatively, bacterial growth was quantified using a qPCR-based method as previously described (Ross and Somssich, 2016). In brief, DNA of bacteria-infiltrated leaves was extracted using the FastDNA^TM^ Spin Kit (MP Biomedicals). Extracted DNA was quantified and adjusted to 8.75 µg/µl to achieve a final concentration of 35 µg DNA in a qPCR reaction. Bacterial DNA was quantified using the levels of the *Pto*-specific *oprF* gene relative to plant *ACTIN2* (*ACT2*) DNA. ΔCt values were calculated by subtracting the Ct value of the target gene from that of *ACT2*. These ΔCt values were considered log_2_ values and used for statistical analysis. Used primers are listed in Supplemental Data Set 9.

### MAP kinase phosphorylation assay

The MAPK phosphorylation assay was performed as previously described (Tsuda et al., 2009). In short, 12-day-old seedlings were treated with 1 µM flg22 or mock for 15 min, frozen in liquid nitrogen and ground with four metal beads in a Retsch MM 400 mixing mill (Retsch, Germany). Then, 150 µl of MAPK extraction buffer (50 mM Tris-HCL [pH 7.5], 5 mM EDTA, 5 mM EGTA, 2 mM DTT, 10 mM NaF, 50 mM ß-glycerolphosphate, 10% glycerol, complete proteinase inhibitor and PhosSTOP phosphatase inhibitor [both from Roche, Germany]) were added to the sample and protein was extracted by centrifugation (4°C, 12,000 rpm). Protein concentration was determined using the Coomassie Protein Assay Kit (ThermoFisher Scientific, USA), and 25 µg of protein were separated by SDS-PAGE for 1 h at 100V. MAPK phosphorylation was detected via immunoblotting using an antiphospho-p44/42 MAPK antibody (dilution 1:5000 in TBST, Cell Signaling Technology, USA) as primary and HRP-conjugated anti-rabbit IgG (1:10000 in TBST, Sigma-Aldrich, USA) as secondary antibody. Luminescence was detected using SuperSignal West Femto Chemiluminescent Reagent (Thermo Fisher Scientific) and a ChemiDoc MP imaging system (Biorad, USA).

### RNA extraction, cDNA synthesis, and RT-qPCR

Seedling samples were ground in 2-mL reaction tubes with four metal beads using a Retsch MM 400 mixing mill (Retsch, Germany). RNA was extracted using peqGOLD TriFast^TM^ with an additional DNA digestion step using DNAse I (Roche, Germany). Further, RNA was precipitated overnight at 4°C in 100% ethanol containing 115 mM Na-Ac (pH 5.2; Sigma Aldrich, Germany) to further clean up and increase RNA yield. RNA quality and quantity was determined using a NanoDrop photometer (Thermo Fisher Scientific). Subsequently, cDNA was synthesized from 4 µg DNAse-treated total RNA using Superscript II or IV Reverse Transcriptase (Thermo Fisher Scientific) according to the manufacturer’s instructions. qPCR was performed on a CFX Connect Real-Time PCR Detection System (Biorad) using EvaGreen (Biotium, USA). The target gene was quantified relative to the expression of *ACTIN2* (*ACT2*) from *A. thaliana* or other Ct values were calculated by subtracting the Ct value of the target gene from that of *ACT2*. These ΔCt values were considered log_2_ values and were further used for statistical analysis. Primers used are listed in Supplemental Data Set 9.

### Statistical analysis

Statistical analysis for the seedling growth inhibition assay, bacterial growth assay and RT-qPCR was performed using a mixed linear model with the function lmer implemented in the lme4 package within the R environment. To meet the assumptions of the mixed linear model, we log-transformed raw data when needed. The following model was fit to the data: measurement_gyr_ ∼GY_gy_ + R_r_ + □_gyr,_ with GY denoting the genotype:treatment interaction effect; R, biological replicate effect; □, residual. The p-values calculated in two-tailed t-tests were corrected for multiple hypothesis testing using the qvalue package when samples were compared with each other in a given figure panel.

### RNA-seq

RNA quality was checked with the Agilent 2100 Bioanalyzer or Caliper LabChip GX device. PolyA enrichment and library preparation were performed with the NEBNext Ultra Directional RNA Library Prep Kit for Illumina (New England Biolabs). Libraries were quantified by fluorometry, immobilized and processed onto a flow cell with a cBot (Illumina), followed by sequencing-by-synthesis with HiSeq v3 chemistry. Library construction and RNA sequencing were performed by the Max Planck-Genome-centre Cologne (http://mpgc.mpipz.mpg.de/home/) with single 100 bp (*A. thaliana* Col-0, *C. rubella*, *C. hirsuta*, and *E. salsugineum*) or 150 bp reads (all other *A. thaliana* accessions) using the Illumina HiSeq2500 or HiSeq3000 platforms, respectively. After quality control, raw sequencing reads were mapped to respective reference genomes (Table 4) using TopHat2 (v2.1.1) with default parameters except for the parameters described in Table 5. The resulting bam files were used to count the number of reads per gene using HtSeq (v 0.6.0) software with default parameters. To exclude biases caused by mapping sequence reads of different *A. thaliana* accessions to the Col-0 genome, we created mapping genome files for each *A. thaliana* accession by correcting the Col-0 reference genome with SNP data available for these accessions. We downloaded the variants table for each accession from the website of the 1001 Genomes Project (intersection_snp_short_indel_vcf V3.1 dataset). The pseudo-genome sequence of each accession was inferred by replacing the reference allele with the corresponding alternative allele using the getGenomeSequence function implemented in AnnotationLiftOver software (https://github.com/baoxingsong/AnnotationLiftOver). Further we created general feature format files (GFF) by projecting the coordinates of the TAIR10 gene annotations onto the coordinates of each accession with the gffCoordinateLiftOver function of AnnotationLiftOver. With these files, we performed a second mapping as described above. The RNA-seq data used in this study are deposited in the National Center for Biotechnology Information Gene Expression Omnibus database (accession no. GSE115991).

**Table 4:** Reference genomes used for RNAseq analysis.

| Species | Reference genome | publication | Source |
| --- | --- | --- | --- |
| *Arabidopsis thaliana* | TAIR 10 | (Lamesch et al., 2012) | Phytozome 10 |
| *Ath accessions* | SNP corrected TAIR10 |  | This study |
| *Capsella rubella* | v1.0 | (Slotte et al., 2013) | Phytozome 10 |
| *Cardamine hirsuta* | v1.0 | (Gan et al., 2016) | http://chi.mpipz.mpg.de/ |
| *Eutrema salsugineum* | v1.0 | (Yang et al., 2013) | Phytozome 10 |

**Table 5:** Tophat2 parameters used for mapping RNAseq reads.

| TopHat2 parameter | Value |
| --- | --- |
| --read mismatches | 10 |
| -- read-gap-length | 10 |
| -- read-edit-dist | 20 |
| --min-anchor-length | 5 |
| --splice-mismatches | 2 |
| --min-intron-length | 30 |
| --max-intron-length | 1000 |
| --max-insertion-length | 20 |
| --max-deletion-length | 20 |
| --max-multihits | 10 |
| --segment-mismatches | 3 |
| --min-coverage-intron | 30 |
| --max-coverage-intron | 10000 |
| --library-type | fr-firststrand |
| --b2 | very sensitive |

The read counts determined by HTSeq were analysed in the R environment (v.3.3.1) using the edgeR (version 3.14.0) and limma (version 3.28.14) packages. Lowly expressed genes were excluded from analysis by filtering out genes with a mean read count below 10 counts per sample. Then, read counts were normalized using TMM normalization embedded in the edge R package and the data were log_2_-transformed using the voom function within the limma package to yield log_2_ counts per million. For individual analysis of Brassicaceae species and *A. thaliana* accession data, a linear model was fit to each gene using the lmFit function of limma with the following terms: S_gyr_ = GY_gy_ + R_r_ +□_gyr_, where S denotes log_2_ expression value, GY, genotype:treatment interaction, and random factors are R, biological replicate; □, residual. For the combined analysis of Brassicaceae species and *A. thaliana* accession data the replicate effect was removed from the linear model resulting in the following terms: S_gy_ = GY_gy_ + □_gy_. For variance shrinkage of calculated p-values, the eBayes function of limma was used. The resulting p-values were corrected for multiple testing by calculating the false discovery rate (or q-value) using the qvalue (v.2.4.2) package.

Normalization and determination of DEGs were performed separately for each Brassicaceae species and each *A. thaliana* accession. To compare expression changes mediated by flg22 between Brassicaceae plants, we used Best Reciprocal BLAST to determine genes which show a 1:1 orthologue with a corresponding *A. thaliana* gene and only kept the genes with 1:1 orthologues in every Brassicaceae species. This resulted in a set of 17,856 1:1 ortholog genes. We restricted the analysis of *A. thaliana* accessions to the same set of 17,856 genes to enable a direct comparison of results obtained from Brassicaceae and *A. thaliana* accession analysis. To directly compare Brassicaceae plants with *A. thaliana* accessions, we further normalized and determined DEGs for all 1-h samples together using the set of 17,856 orthologous genes. This approach enabled us to compare basal expression levels between Brassicaceae and *A. thaliana* accessions.

The R packages and software used for further analysis of the sequencing data are listed in Table 6. Heatmaps and k-mean clustering of DEGs were generated using the Genesis software with default parameters.

**Table 6:** Software and packages used in this study.

| Software/Package | Version | Citation | Use |
| --- | --- | --- | --- |
| AME | 4.12.0 | (McLeay and Bailey, 2010) | TF motif enrichment |
| BinGO | 3.0.3 | (Maere et al., 2005) | GO enrichment |
| Clustal Omega | 1.2.4 | (Sievers et al., 2011) | Multiple sequence alignment |
| Cytoscape | 3.3.0 | (Shannon et al., 2003) | Run BinGO |
| EdgeR | 3.14.0 | (Robinson et al., 2010) | Analysing DEGs |
| Genevestigator |  | (Hruz et al., 2008) | public transcriptome data |
| Genesis | 1.7.7 | (Sturn et al., 2002) | Heatmaps, clustering |
| Htseq | 0.6.0 | (Anders et al., 2015) | Count RNAseq reads |
| limma | 3.28.14 | (Ritchie et al., 2015) | Analysing DEGs |
| MixOmics | 6.0 | (Rohart et al., 2017) | PCA |
| TopHat | 2.1.1 | (Trapnell et al., 2009) | Map RNAseq reads |
| VennDiagramm | 1.6.17 | (Chen and Boutros, 2011) | Venn Diagramms |

The expression clusters of DEGs determined for the combined RNAseq analysis of *A. thaliana* accessions together with Brassicaceae species were investigated for enrichment of GO terms corresponding to biological processes using the BinGO plugin within the Cytoscape environment. GO term enrichment was calculated using a hypergeometric test followed by Benjamini and Hochberg False Discovery Rate correction implemented in the BinGO plugin. The whole genome annotation was used as a background.

Known TF motifs enriched in individual expression clusters of DEGs determined fin the combined RNAseq analysis of *A. thaliana* accessions together with Brassicaceae species were identified using the AME tool within the MEME suite. For this purpose, 5′-gene regulatory regions (500 bp upstream of the transcription start site) were extracted for each tested Brassicaceae species. Enrichment of TF motifs was determined in each of the 15 k-means clusters for all tested Brassicaceae species using the 5′□regulatory-regions of all expressed genes having clear 1:1 orthologues (16,100 genes) as a background. Known TF motifs were retrieved from the JASPAR CORE (2018) plants database that is implemented in AME.

To compare amino acid sequence conservation with expression variation, all amino acid sequences of expressed genes with 1:1 orthologues in all species were extracted for each Brassicaceae species. The sequences were aligned using Clustal Omega and percent identity matrices were extracted. The amino acid sequence identity output of Clustal Omega was used to calculate the mean amino acid identity across *C. rubella*, *C. hirsuta* and *E. salsugineum* compared to *A. thaliana* as a proxy of sequence conservation. The mean amino acid sequence identities were subsequently plotted against the SD/mean of flg22-expression changes across all four Brassicaceae species, which served as a proxy for expression variation among the tested Brassicaceae species. Similarly, the mean amino acid sequence identity was also plotted against the SD/mean of the normalized expression value in control samples. In addition, pairwise amino acid sequence identities between *A. thaliana* and each Brassicaceae species were plotted against the absolute difference in flg22-induced expression changes between the compared species. This analysis was performed for all expressed genes or only for DEGs.

### Phylogeny analysis

Various dates have been reported for the divergence of Brassicaceae species (Franzke et al., 2016). For instance, Beilstein et al. dated the Brassicaceae crown node age to 54 million years ago (Mya) whereas more recent publications dated this event 31 to 37 Mya ago (Beilstein et al., 2010; Edger et al., 2015; Hohmann et al., 2015; Huang et al., 2016; Franzke et al., 2016). Therefore, in this study, we used TIMETREE (www.timetree.org) which synthesizes divergence times based on the available literature to estimate the timescale of Brassicaceae species evolution (Hedges et al., 2015). Phylogenetic trees were retrieved from timetree.org based on divergence time estimates from 15 studies (Arakaki et al., 2011; Artyukova et al., 2014; Beilstein et al., 2010; Couvreur et al., 2010; Franzke et al., 2009; Heenan et al., 2002; Hermant et al., 2012; Hohmann et al., 2015; Huang et al., 2016; Koch et al., 2000; Mandáková et al., 2010; Naumann et al., 2013; Parkinson et al., 2005; Vanneste et al., 2014; Yue et al., 2009).

### Genevestigator analysis

The following datasets were used for Genevestigator analysis: AT-00106 (*Pto* DC3000); AT-00110 (ABA or MeJA); AT-00113 (SA); AT-00147 (*B. cinerea*); AT-00253 (flg22 or OG); AT-00493 (hypoxia); AT-00553 (*Hyaloperonospora arabidopsidis*); AT-00560 (drought); AT-00597 (Pep2 and elf18); AT-00645 (heatstress)

### SA analysis

SA levels were analysed as described previously with an ultra-high performance liquid chromatography/Q-Exactive™ system (Thermo Fisher Scientific) using an ODS column (AQUITY UPLC BEH C18, 1.7 μm, 2.1 × 100 mm; Waters) (Kojima and Sakakibara, 2012; Kojima et al., 2009)(Yasuda et al., 2016).

### Secondary metabolite extraction, acquisition and processing of data

Control and flg22-treated seedlings were collected and extracted as described before (Bednarek et al., 2011). The obtained extracts were subjected to LC-MS analyses performed using the Acquity UPLC system (Waters, USA) hyphenated to a micrOToF-Q mass spectrometer (Bruker Daltonics, Germany). Chromatographic separations were carried out on a BEH C18 column (2.1×150 mm, 1.7 μm particle size) at 22°C with a mobile phase flow rate of 0.35 ml/min. The elution was conducted using water containing 0.1% formic acid (Sigma Aldrich, Germany) (solvent A) and acetonitrile (VWR Chemicals; France) containing 1.9% of water and 0.1% of formic acid (solvent B) in the following gradient: 0–10 min from 0% to 25% B, 10–15 min to 30% B, 20–24 min maintained at 100% B, and up to 24.5 min the system was returned to starting conditions and re-equilibrated for 8 min. Calibration of the spectrometer with sodium formate salt clusters was done prior to each analysis. MS was operated using the following settings: ion source voltage of -4.5 kV or 4.5 kV, nebulization of nitrogen at a pressure of 1.2 bar and a gas flow rate of 8 l/min. Ion source temperature was 220°C. The spectra were scanned in positive and negative ion mode at the range of 50–1000 *m/z* at a resolution higher than 15,000 FWHM (full width at half maximum). Data acquisition was supervised by HyStar 3.2 software (Bruker Daltonics, Germany).

Obtained LC-MS data were converted to the *mzXML* format by MSConvert Version: 3.0.11781 tool available in Proteowizard software prior to further processing by MZmine 2.31 software (Pluskal et al., 2010). Data from each experiment were processed separately for negative and positive ionization. In first step, lists of masses were generated by the mass detector module in each scan in the raw data files. Then, chromatograms for each mass detected continuously over the scans were built using a chromatogram builder algorithm. These chromatograms were deconvoluted by the deconvolution module using the wavelets algorithm based on Bioconductor’s XCMS package for R (Tautenhahn et al., 2008). An isotopic peaks grouper was used for isotope elimination followed by adduct and complex searching. Deviation of retention times between peak lists was reduced by a retention time normalizer. Such transformed peaks were aligned in all samples through a match score by a join aligner module. The resulting peaks list was completed by supplemental peak detection with a peak finder algorithm prior to missing value imputation (gap filling). The generated data table was subsequently exported in *csv* format for further statistical analysis.

Observations equal to zero (below the detection level) were substituted by half of the minimum non-zero observation for each metabolite. Then observations were transformed by log_2_(10^3^x). Two-way analysis of variance (ANOVA) was done with experiment as a block (random effects) and treatment, species as 2 fixed factors; analysis was done together for positive and negative ionization. Differentially Accumulated Metabolite (DAM) was indicated for each species if all of three conditions hold: i) treatment effect or interaction treatment x species was significant with q-value < 0.05 (fdr – false discovery rate, (Benjamini and Hochberg, 1995)), ii) individual tests for each fungi p < 0.05 (significant test for the difference between treatment and control was done for each species), iii) |fold change|>1.5, where fold change is flg22 treatment/control. Statistical analysis was performed in Genstat 19. Visualizations including barplot, PCA, heatmap and Venn diagram were created in R.

## Acknowledgements

We thank the members of the Tsuda lab for critical reading of the manuscript, the Max-Planck Genome Centre for sequencing support, and Neysan Donnelly and Tsuda lab members for critical comments on the manuscript. This work was supported by the Huazhong Agricultural University Scientific & Technological Self-innovation Foundation, the Max Planck Society, a German Research Foundation grant (SPP2125) (to K.T.), IMPRS Cologne (T.M.W) and National Science Center OPUS (UMO-2015/17/B/NZ1/00871) (to P.B) and PRELUDIUM (UMO-2013/09/N/NZ2/02080) (to K.K) grants.

## Author contributions

T.M.W., S.A., P.S.-L., P.B. and K.T. designed the research. T.M.W., F.E., S.A., P.A., A.P., and K.K. performed research. H.S., X.G., M.T., H.S., and R.G.-O. contributed new reagent/analytic tools. T.M.W., S. A., F.E., A.P., B.S., E.D., A.S., K.F., S.L., P.B., and K.T. analyzed data. T.M.W., F.E., and K.T. wrote the paper with input from all authors.

## Competing interests

The authors declare no competing interest.

---

**Figure S1.**
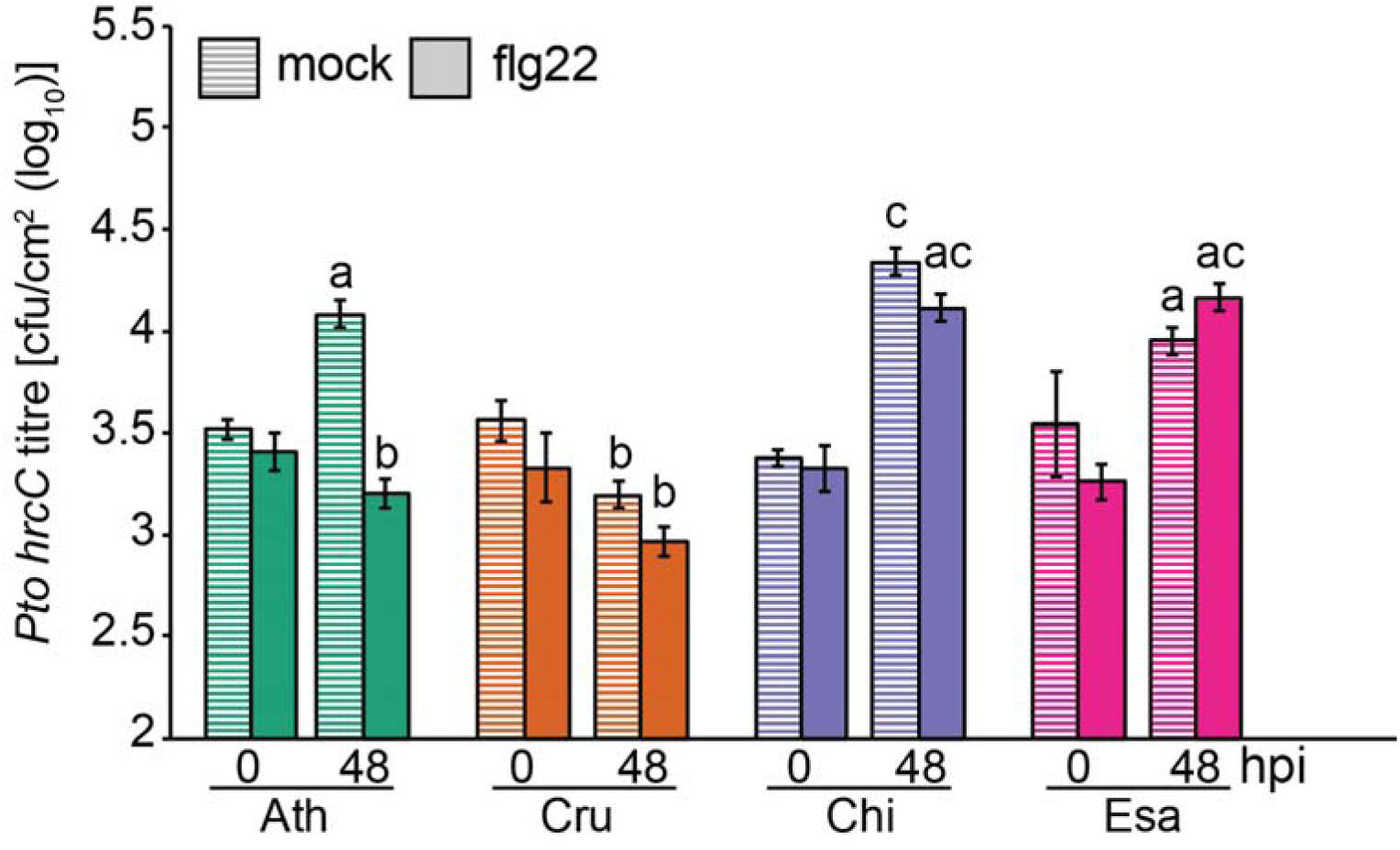
Effects of flg22 on *Pto hrcC* growth in Brassicaceae species. (Supports Figure 1) Leaves of 5-week-old plants were syringe-infiltrated with mock or 1 μM flg22 24 h prior to infiltration with *Pto hrcC* (OD_600_ = 0.001). The bacterial titre was determined 0 and 48 h after bacterial infiltration. Bars represent means and SEs from 2 independent experiments each with 12 biological replicates (n = 24). Different letters indicate statistically significant differences (mixed linear model, adjusted p < 0.01). Ath, *A. thaliana* (Col-0); Cru, *C. rubella* (N22697); Chi, *C. hirsuta* (Oxford); Esa, *E. salsugineum* (Shandong).

**Figure S2.**
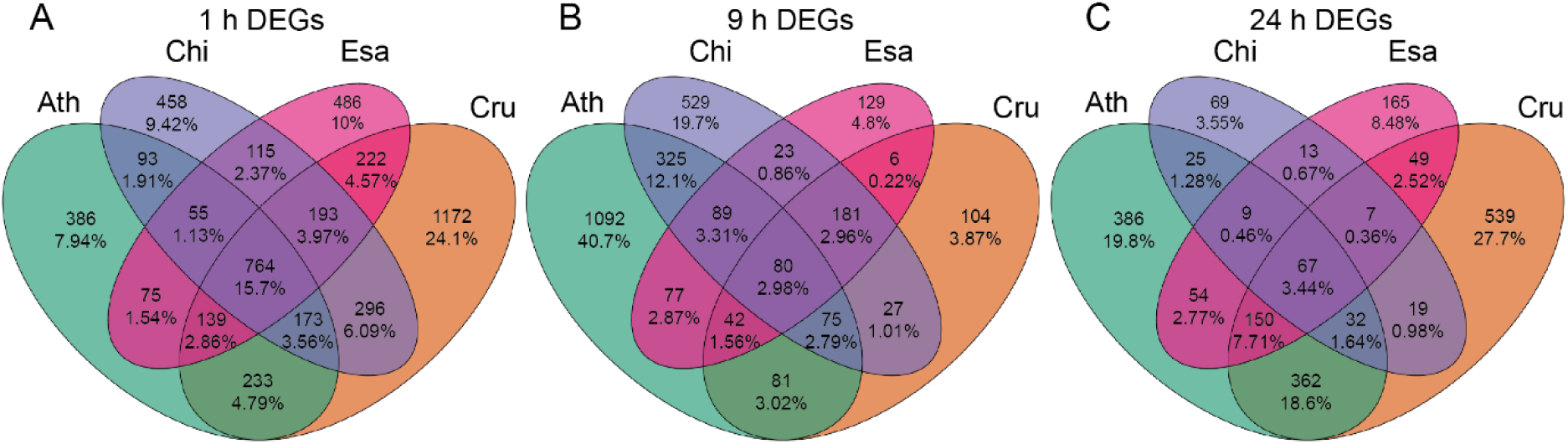
Overlap of DEGs at each time point. (Supports Figure 2) Venn diagrams showing shared and specific DEGs between species at 1 h (**A**), 9 h (**B**) and 24 h (**C**) after flg22-treatment. All DEGs differentially expressed in at least species at the respective time points were used. Ath, *A. thaliana* (Col-0); Cru, *C. rubella* (N22697); Chi, *C. hirsuta* (Oxford); Esa, *E. salsugineum* (Shandong).

**Figure S3.**
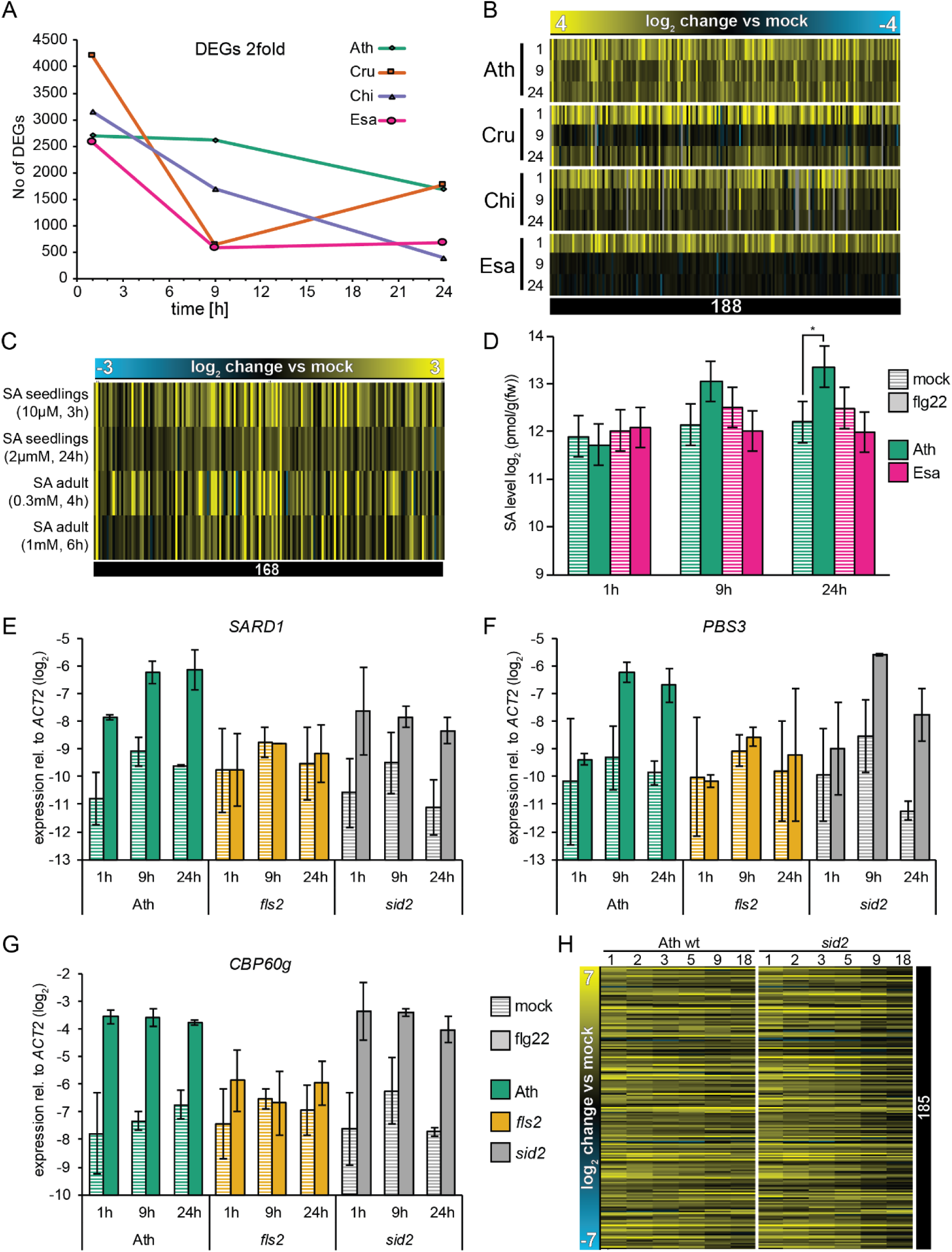
*SID2*-dependent SA accumulation is not required for sustained transcriptome responses in *A. thaliana*. (Supports Figure 3) (**A**) The temporal dynamics of the transcriptional response to flg22 differs in Brassicaceae species. The numbers of DEGs (q-value < 0.01; |log2 fold change| > 1) at each time point in each species are plotted. (**B**) Heatmap visualizing 188 genes induced at 1 hpt in Ath and Esa (log2 induction > 0.6) with sustained induction in Ath (log2 induction > 0.6 at 9 and 24 hpt) but transient induction in Esa (log2 induction < 0.5 at 9 and 24 hpt). (**C**) Most of the 188 genes (missing genes are due to missing probes on microarrays of public datasets) are responsive to SA in publically available expression data of *A. thaliana* (Genevestigator). (**D**) Free salicylic acid (SA) levels of 12-day-old seedlings were determined using HPLC-MS at the indicated time-points after mock or 1 μM flg22 treatment. Bars represent the means ±SE from 3 independent experiments. Asterisks indicate significant difference to mock (mixed linear model followed by Student’s t-test; *, p < 0.05). (**E – G**) 12-day-old seedlings of *A. thaliana* Col-0 wild type, *fls2*, and *sid2* were treated with mock or 1 μM flg22 for 1, 9, or 24 h. Expression of 3 marker genes extracted from the heatmap in Figure 4B, namely *SARD1* (**E**), *PBS3* (**F**) and *CBP60g* (**G**) was quantified using RT-qPCR. Bars represent the means ±SD from 2 independent experiments. Ath, *A. thaliana* (Col-0); Cru, *C. rubella* (N22697); Chi, *C. hirsuta* (Oxford); Esa, *E. salsugineum* (Shandong). (**H**) The 188 genes in Figure 4B showing transient induction in Esa were analyzed for their expression induction in 31 to 32-day-old Col-0 and *sid2* leaves at the indicated time points compared to 0 h after 1 μM flg22 treatment (Hillmer et al., 2017).

**Figure S4.**
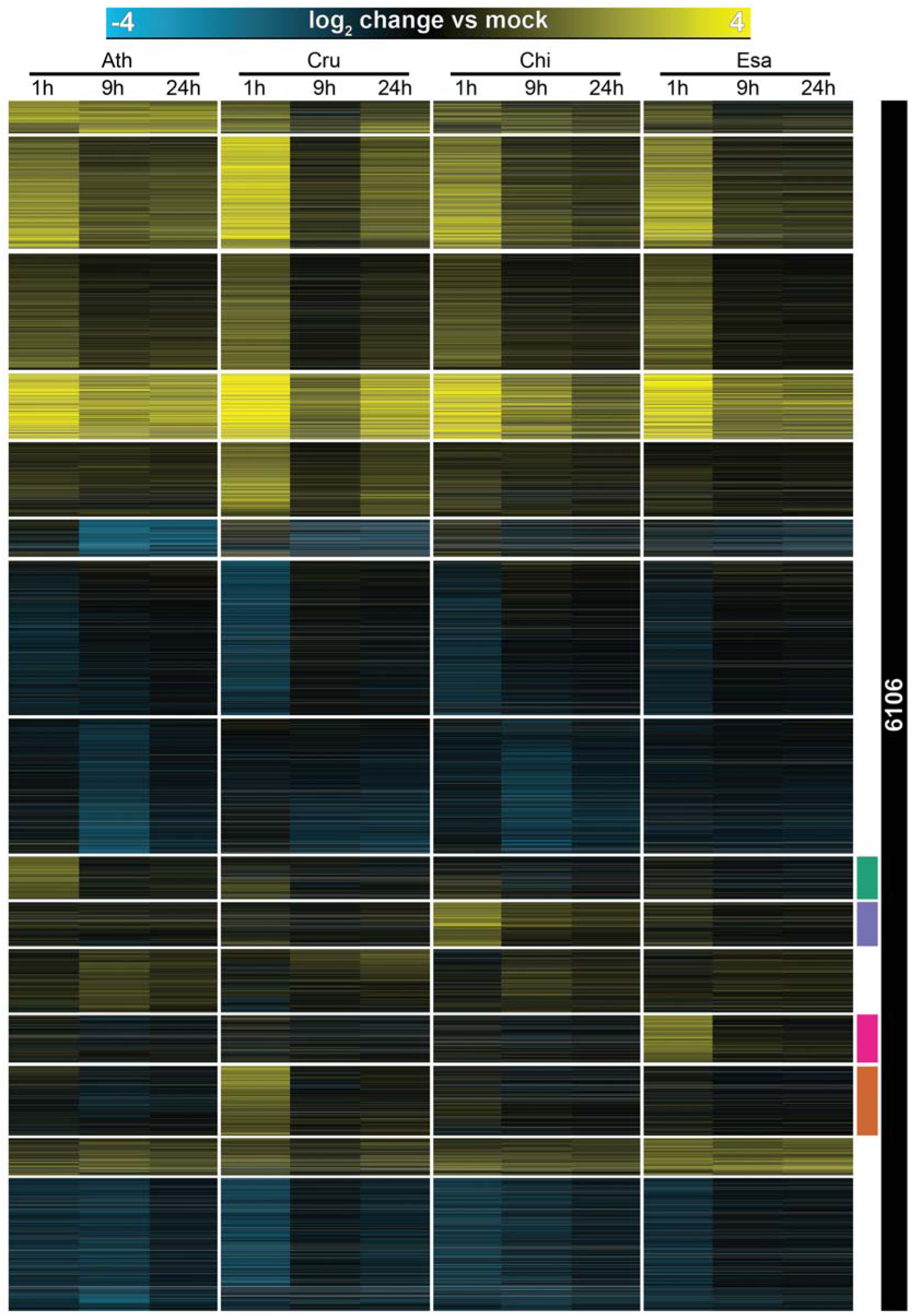
Heatmap for all DEGs in Brassicaceae species after flg22 treatment. (Supports Figure 3) Heatmap of log_2_ fold changes for all 6,106 DEGs among all Brassicaceae species generated using k-means clustering (k = 15). All DEGs which are differentially expressed at least at 1 time point in 1 species were used. Species-specific expression clusters shown in Figure 3C are indicated by coloured bars [Ath (green), Cru (orange), Chi (purple), and Esa (magenta)] on the right side. Ath, *A. thaliana* (Col-0); Cru, *C. rubella* (N22697); Chi, *C. hirsuta* (Oxford); Esa, *E. salsugineum* (Shandong).

**Figure S5.**
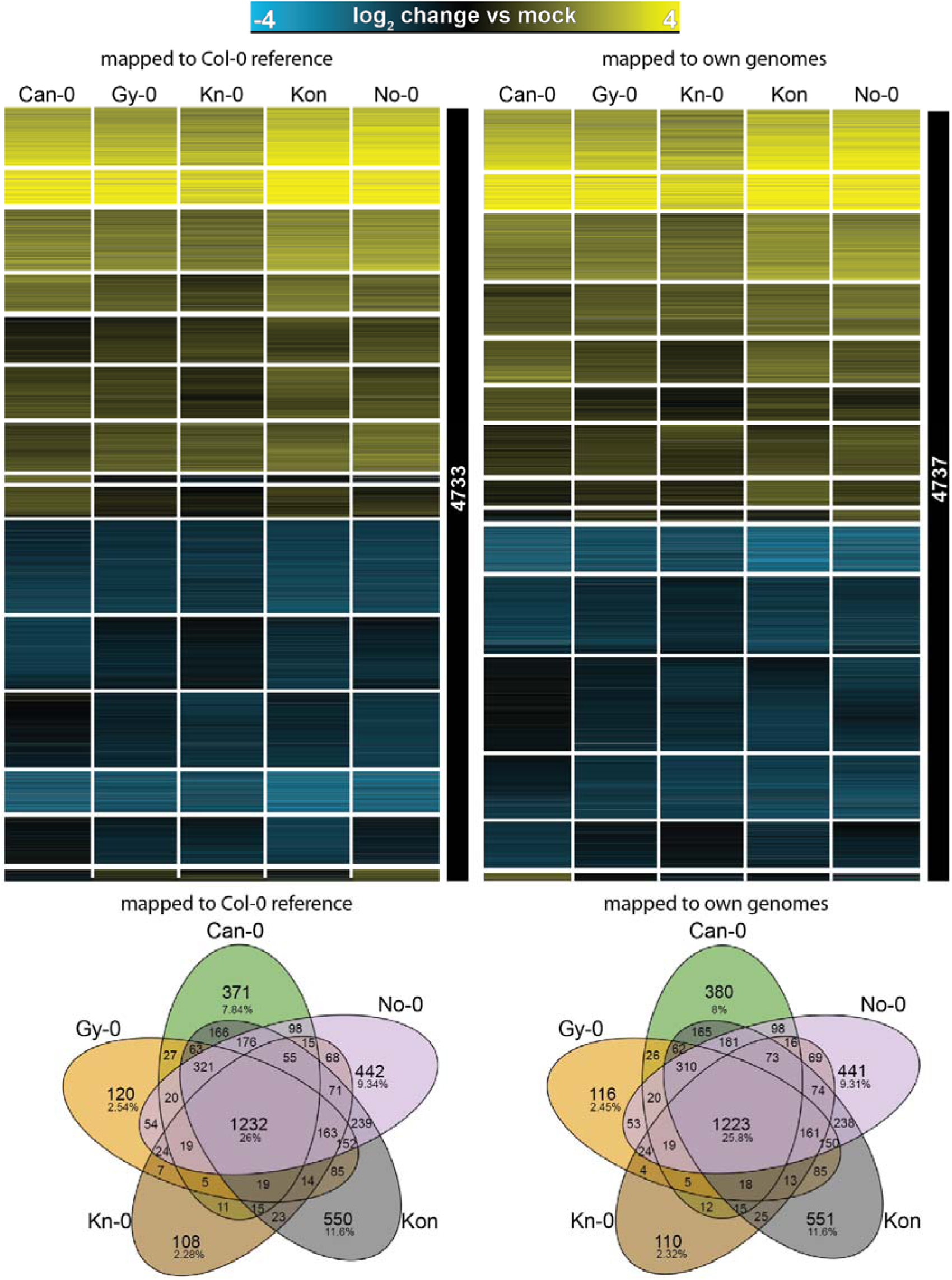
Comparison of two different mapping approaches for *A. thaliana* accession RNA-seq reads. (Supports Figure 4) RNA-seq reads were mapped to the Col-0 (TAIR10) reference genome (left) or to individual *A. thaliana* accession genomes generated in this study using SNP data (right). Heatmap of DEGs in at least 1 accession clustered by k-means (k = 15). Log_2_ expression changes compared to mock are shown.

**Figure S6.**
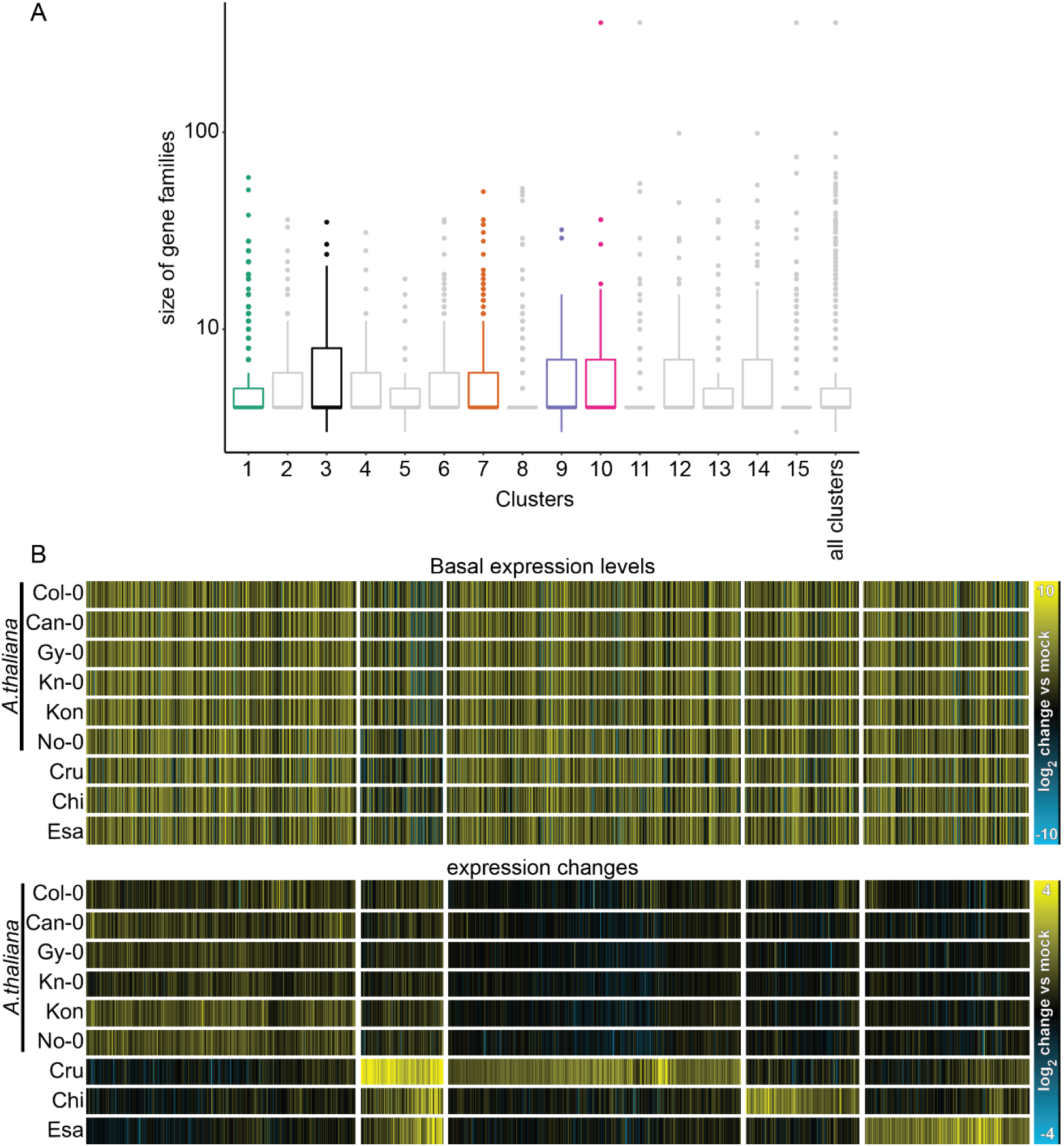
Gene family size and basal gene expression levels do not explain species-specific expression signatures. (Supports Figure 5) (**A**) The sum of the number of genes in gene families among the 4 tested Brassicaceae species are plotted at log_10_ scale for each of the 15 clusters obtained by k-means clustering of all 5,961 genes that are DEGs in at least species or accession (See Figure 5). Species-specific clusters are highlighted by colours [Ath (green), Cru (orange), Chi (purple) and Esa (magenta)]. Cru, *C. rubella* (N22697); Chi, *C. hirsuta* (Oxford); Esa, *E. salsugineum* (Shandong)]. (**B**) Basal (mock condition) expression levels (normalized and log_2_-transformed counts per million) of genes showing species-specific expression signatures are shown in the upper heatmap. Log_2_ expression changes after flg22 treatment are shown in the bottom heatmap.

**Figure S7.**
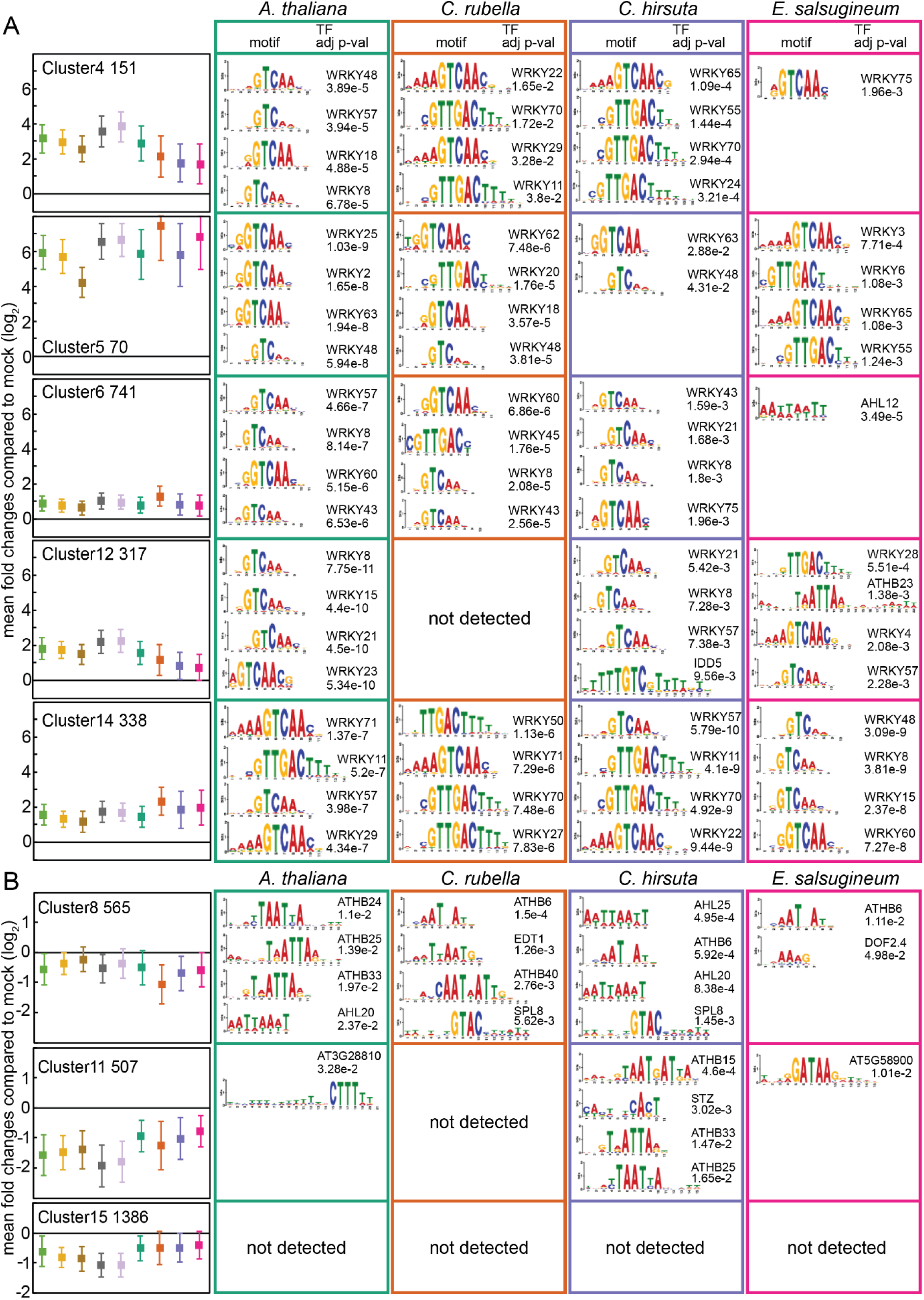
Enrichment of TF-motifs within the 5′-regulatory regions of DEG clusters. (Supports Figure 7) The 500 bp upstream sequences of the transcription start sites of the genes in the individual clusters were tested for enrichment of known TF binding motifs. Names of transcription factors, sequence logos and adjusted p-values (up to the top 4) of motifs are shown for each Brassicaceae species. The names of clusters, the number of DEGs, and mean log_2_ fold changes ±SD compared to mock are shown on the left side. For the complete list of all enriched TF binding motifs, please see Supplemental Data Set 6. Ath, *A. thaliana* (Col-0); Cru, *C. rubella* (N22697); Chi, *C. hirsuta* (Oxford); Esa, *E. salsugineum* (Shandong). A: Commonly induced clusters. B: Commonly downregulated clusters.

**Figure S8.**
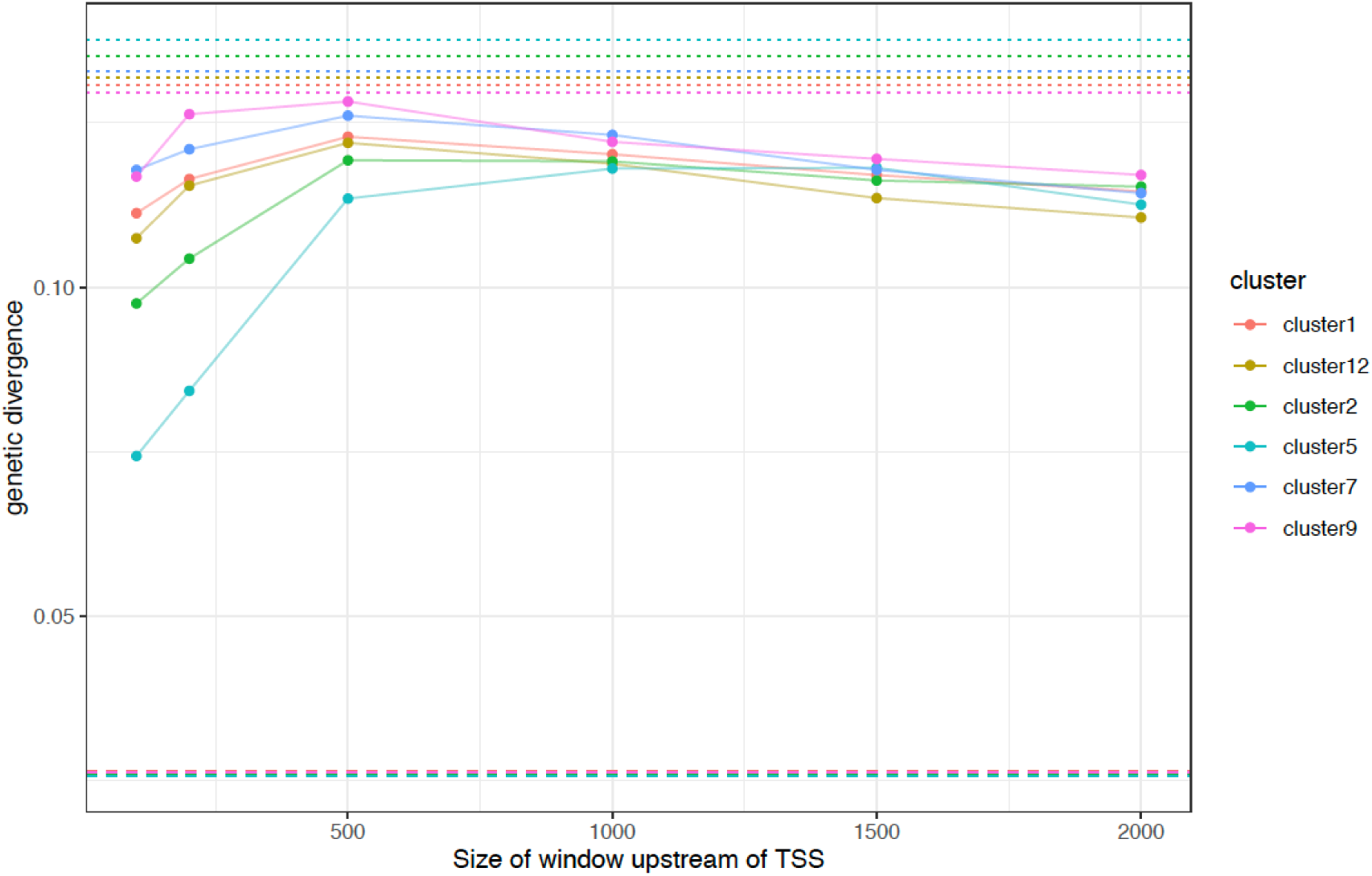
Genetic divergence between *A. thaliana* and *A. lyrata* for upstream, synonymous, and non-synonymous sites. (Supports Figure 8) The y-axis shows the proportion of sites that are different between the reference sequences of *A. thaliana* and *A. lyrata*. The x-axis indicates the size of the window upstream of the SFS used to calculate the “upstream” genetic divergence (solid lines). The dashed and dotted lines represent the non-synonymous and synonymous genetic divergences, respectively.

**Figure S9.**
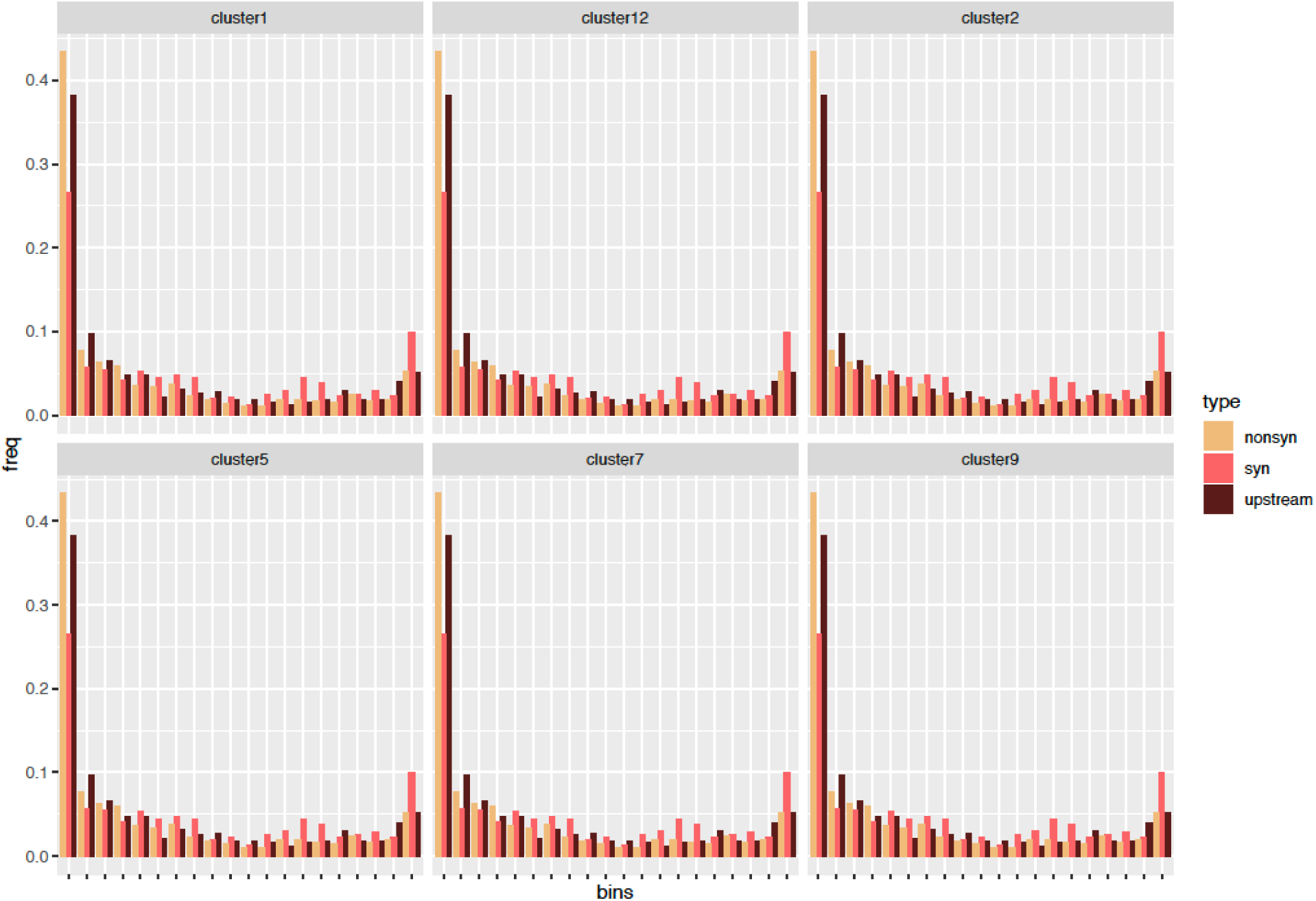
Distribution of allele frequencies (SFS) at upstream and coding sequences in *A. thaliana*, in 6 clusters. (Supports Figure 8) The x-axis represents discretized frequency classes for all alleles found in our population sample from the 1001 genome project (n=45). Variation is partitioned by expression cluster (6) and functional class (1000 bp upstream of start codon, nonsynonymous and synonymous sites). The y-axis represents the proportion of alleles falling in each frequency class.

**Figure S10.**
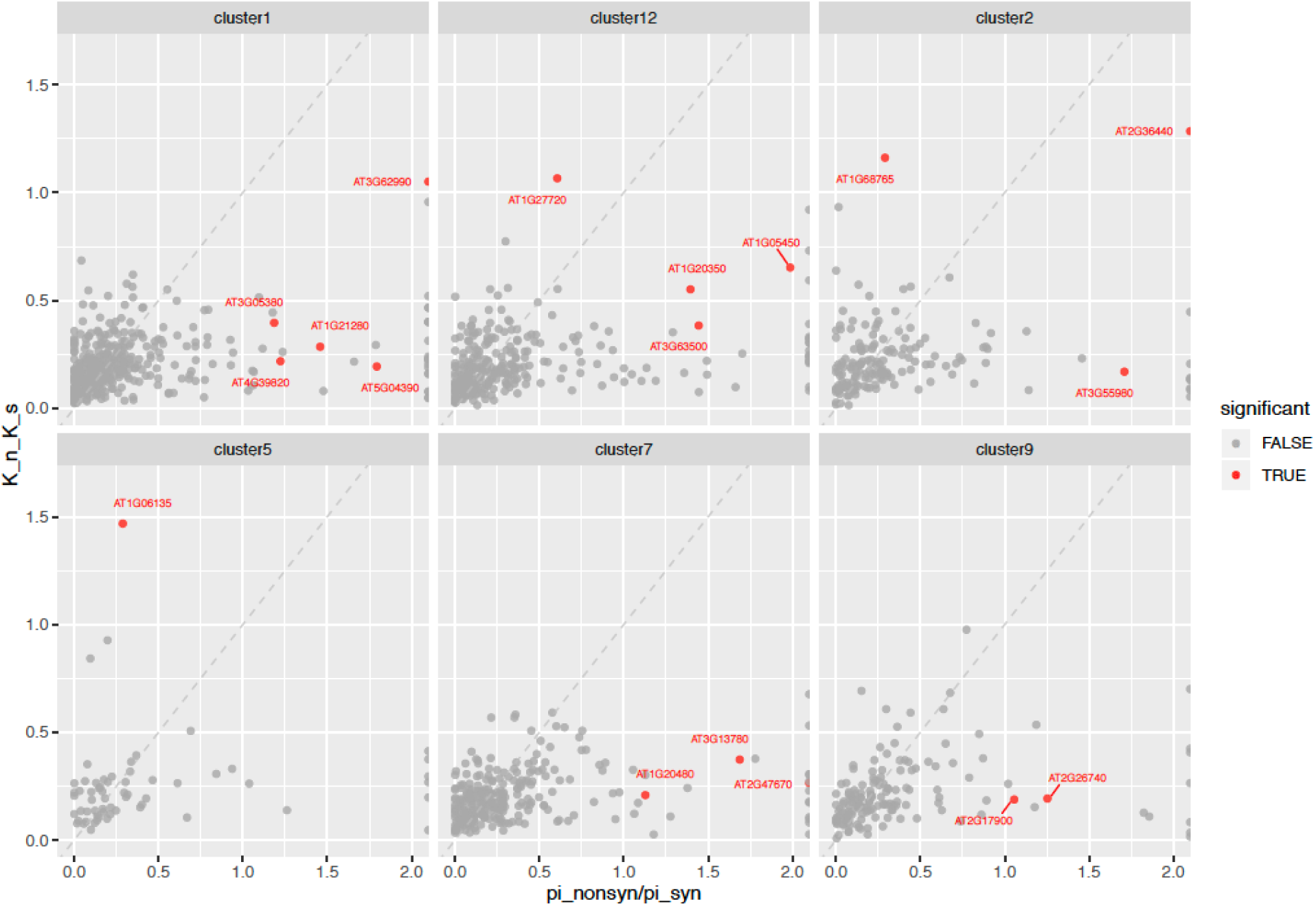
Long-term balancing selection has been identified as an important process shaping the genetic variation of immunity related genes in plant. (Supports Figure 8) One of the signatures of balancing selection in polymorphism and divergence data is to increase the ratio of non-synonymous to synonymous divergence and polymorphism (Ka/Ks and pi_a/pi_s, respectively) with which the values for each gene (coding sequence) partitioned by cluster. Genes with a value larger than 1 for one of those ratios and that have at least 4 non-synonymous polymorphic sites have been highlighted as outlier.

## Notes

### Competing Interest Statement

The authors have declared no competing interest.

### Summary of Updates

Supplemental Data Sets 1 to 9 have been added. Other parts remain the same as the previous version.

